# IL-17A-mediated alterations in gut microbiota composition, particularly changes in Prevotella abundance, impact Treg function in CNS Autoimmunity

**DOI:** 10.1101/2022.04.22.489206

**Authors:** Shailesh K. Shahi, Sudeep Ghimire, Samantha N. Jensen, Peter Lehman, Allison G Rux, Ti-Ara Turner, Nicholas Borcherding, Katherine N. Gibson-Corley, Sukirth M. Ganesan, Nitin J. Karandikar, Ashutosh K. Mangalam

## Abstract

A disrupted equilibrium among gut microbiota, IL-17A-producing CD4 T-cells (Th17), and regulatory CD4 T-cells (Treg) have been linked with the pathobiology of multiple sclerosis (MS). While gut microbiota can regulate both Treg and Th17 cells, the impact of IL-17A on this gut-immune connection remains unclear. Utilizing HLA-DR3 transgenic mouse model of MS, we show that IL-17A deficiency (HLA-DR3.IL17A^-/-^) resulted in milder disease characterized by increased Tregs and expansion of Treg-promoting gut microbes, including *Prevotella*. Cohousing HLA-DR3 mice with HLA-DR3.IL17A^-/-^ transferred the milder disease phenotype and associated microbiota changes to DR3 mice, highlighting the dominant role of gut microbiota in Treg induction and disease amelioration. DR3.IL17A^-/-^ mice also showed a higher abundance of functional pathways linked with short-chain fatty acid synthesis and elevated IL-10 in dendritic cells. Enrichment of the Treg-promoting PPAR signaling pathway expression in the colon of HLA-DR3.IL17A^-/-^ mice and following *Prevotella* administration in HLA-DR3 mice underscores the importance of gut microbiota in IL-17A-mediated immune regulation. Thus, our study uncovers a previously unappreciated role for IL-17A in shaping gut microbiota and immune regulation, with far-reaching implications for MS treatment.

**One-Sentence Summary:** IL-17A modulates Treg and gut microbiota to control EAE

## Main text

Multiple sclerosis (MS) is a chronic, inflammatory, neurodegenerative disease of the central nervous system (CNS), in which CD4 T helper (Th) cells of Th1 and/or Th17 phenotype have been shown to play an important role in pathobiology of the disease. IL-17A is a signature pro-inflammatory cytokine of the CD4 T helper 17 (Th17) subset that has emerged as being critical to the pathobiology of MS (1, 2). IL-17A plays a crucial role in resolving extracellular bacterial and fungal infections, by recruiting immune cells, specifically neutrophils, to the site of infection and inducing pro-inflammatory mediators from immune cells and fibroblasts, epithelial cells, and endothelial cells (3). However, an aberrant IL-17A response can lead to unresolved inflammation and tissue pathology, as observed in multiple autoimmune diseases, including MS (4, 5, 6, 7). Thus, although IL-17A is considered a major contributor to the pathogenesis of multiple autoimmune diseases, its precise role in the pathobiology of MS is still being explored. Th17 cells are regulated by CD4^+^CD25^+^FoxP3^+^ regulatory T cells (Treg) and the balance between Treg/Th17 is shown to shift toward the Th17 phenotype in inflammatory diseases such as MS (8). Additionally, gut bacteria help maintain immune homeostasis by regulating the balance between CD4^+^CD25^+^FoxP3+ regulatory T cells (Tregs) and pro-inflammatory Th1/Th17 cells (9). The importance of gut microbiota in regulating the development of Treg and Th17 is highlighted by studies performed in germ-free mice where an absence of or reduced Treg and Th17 cells is evident. However, transplant of single bacterium such as segmented filamentous bacterium can restore Th17 (10) and a mixture of chloroform-resistant *Clostridium* (11) can induce/restore Treg population. These studies highlight the important role of gut microbiota in the regulation of the Th17 and Treg population and their role in the pathobiology of MS/EAE. However, whether IL-17A can regulate gut microbiota and/or Treg cells is unknown.

In addition to IL-17A, HLA class II genes, specifically HLA-DR2 and -DR3 alleles, are linked as the strongest genetic factors predisposing individuals to MS (12, 13, 14, 15, 16). We have previously utilized HLA-class II transgenic mice to validate the significance of HLA-class II molecules in the pathogenesis of MS (17, 18, 19, 20, 21). Additionally, we showed that loss of IFNγ was associated with increased IL-17A levels and severe disease in an HLA-class II transgenic mouse model of experimental autoimmune encephalomyelitis (EAE) (22). Collectively these data suggest that IL-17A might be responsible for severe disease in the absence of IFNγ, however it is unclear whether IL-17A is essential for disease induction or severity or both.

To elucidate the role of IL-17A in the pathogenesis of MS, we generated HLA-DR3 (DR3) class II transgenic mice lacking IL-17A (DR3.IL17A^-/-^). Mice that express HLA-DR3 were chosen over other EAE models as our prior study suggested a role of IL-17A in disease severity in the absence of IFNγ (23). Unexpectedly, DR3.IL17A^-/-^ transgenic mice were not resistant to EAE and developed EAE disease with similar disease incidence and onset. However, disease was milder compared to IL-17A-sufficient HLA-DR3 transgenic mice. In the absence of IL-17A, CD4^+^CD25^+^FoxP3^+^ Treg was expanded, in part due to enhanced growth of gut bacteria with the ability to induce Treg. Fecal transplant/cohousing studies confirmed the importance of gut bacteria in modulating IL-17A-mediated disease. IL-17A-sufficient mice that received fecal transplants and cohoused with IL-17A-deficient mice had lower EAE severity compared to DR3 transgenic mice. Mice lacking IL-17A (HLA-DR3.IL17A-/-) exhibited an enriched Treg-promoting PPAR signaling pathway, suggesting a link between gut bacteria and immune regulation. This same pathway showed increased expression after *Prevotella copri* administration in HLA-DR3 mice, further emphasizing the role of gut microbiota in influencing IL-17A-mediated immune responses. Finally, DR3.IL17A^-/-^ transgenic mice showed enriched bacterial pathways for SCFA production and increased IL-10 from dendritic cells, suggesting an anti-inflammatory environment due to IL-17A deficiency. Thus, our study demonstrates that IL-17A regulates Treg levels by modulating the gut microbiota, which alters immune homeostasis and inflammation to promote EAE.

## Materials and methods

### Study design

The study utilized female HLA-DR3, HLA-DR3.IL17A^-/-^ transgenic mice, DR3.IL17A^-/-^IFNγ^-/-^ transgenic mice and C57BL/6J mice to investigate the contributions of IL-17A to EAE. The disease was induced by immunizing mice with the PLP_91-110_ peptide in HLA transgenic mice. C57BL/6 mice were immunized with MOG_35−55_ peptide. The number of sample sizes are indicated in the Figure captions. The disease induction and scoring of the EAE was performed as described below and elsewhere (24). No data, including outliers, were excluded from the analysis. Experiments were performed at least twice with some exceptions as indicated in the Figure legends.

### Mouse procurement, disease induction, and scoring

HLA-DR3 (DRA1*0101, DRB1*0301) transgenic mice on the B6/129 background have been characterized previously (Das et al., 2000; Mangalam et al., 2009). These mice lack endogenous murine major histocompatibility complex (MHC) class II genes (AE^-/-^) and express alleles DRA1*0101, DRB1*0301, as described previously (Das et al., 2000; Mangalam et al., 2009). The HLA-DR3.IL17A^-/-^ mice (referred to as DR3.IL17A^-/-^) that were used in this study were generated by crossing HLA-DR3 (referred to as DR3) and AE^-/-^.IL17A^-/-^ mice and validated using genotyping. HLA-DR3 Transgenic mice were mated with IFNγ-deficient mice on B6 background (Jackson Laboratory) to generate HLA-DRB1*0301.AE^−/−^.IFNγ^−/−^. DR3.IL17A^-/-^IFNγ^-/-^ transgenic mice were generated by crossing HLA-DR3.IL17^-/-^ and HLA-DR3. IFNγ^-/-^ mice and validated using genotyping. C57BL/6 mice were also utilized in this study and were purchased from the Jackson Laboratories (Bar Harbor, ME). Female DR3, DR3.IL17A^-/-^, DR3.IL17A^-/-^IFNγ^-/-^, and C57BL/6 mice (8-10 weeks of age) were used in this study. Mice were bred and maintained in the University of Iowa animal facility in accordance with NIH and institutional guidelines. All experiments were approved by the Institutional Animal Care and Use Committee at the University of Iowa.

To induce EAE in mice, DR3, DR3.IL17A^-/-^, and DR3.IL17A^-/-^IFNγ^-/-^ transgenic mice (8 to 12 weeks old) were immunized subcutaneously in both flanks with 50 µg of PLP_91–110_ (YTTGAVRQIFG DYKTTICGK) (GenScript, Piscataway NJ, USA) that was emulsified in CFA containing 200 μg/mouse *Mycobacterium tuberculosis* H37Ra (Becton, Dickinson and Company, Sparks, MD, USA). 80 ng of pertussis toxin (PTX) (Sigma Chemicals, St. Louis, MO; 100 ng) was given intraperitoneal at day 0 and 2 post immunization. C57BL/6 mice were immunized subcutaneously in both flanks with MOG_35-55_ CFA/PTX, as just described for immunization with PLP (25). Mice were observed daily for disease symptoms and EAE was scored with the following scoring system, as described previously (24): normal, 0; loss of tail tone, 1; hind limb weakness, 2; hind limb paralysis, 3; hind limb paralysis and forelimb paralysis or weakness, 4; and morbidity/death, 5.

### Pathology

Brains and spinal cords were collected from DR3, and DR3.IL17A^-/-^ transgenic mice at day 30 post immunization. Histological analysis was performed to assess inflammation and demyelination, as described previously (22, 26). Briefly, DR3 and DR3.IL17A^-/-^ transgenic mice were perfused with approximately 50 ml of 10% neutral buffered formalin via intracardiac puncture. The calvaria and spinal columns were collected, and immersion fixed in 10 % neutral buffered formalin (10% BFA) for 24-48 hours followed by decalcification in EDTA for approximately 24 hours. Decalcified tissues were cut into 2 mm coronal blocks, embedded in paraffin and routinely processed. The slides were cut at 4-5 mm thickness and stained with hematoxylin and eosin. HE-stained slides including the brainstem, cortex, corpus callosum, cerebellum, hippocampus, and striatum were examined and scored by a board-certified veterinary pathologist as described previously (22, 26).

### Flow cytometry

Peripheral blood mononuclear cells (PBMCs) were isolated from naïve DR3 and DR3.IL17A^-/-^ transgenic mice using Histopaque-1077 (Millipore Sigma, USA) density centrifugation method as per instructions from the manufacturer (Sigma-Aldrich, St. Louis, USA). Briefly, 200 µl of blood was collected in an EDTA-containing Eppendorf tube by retroorbital bleeding. Blood was diluted with 800 ml of phosphate-buffered saline (PBS; Gibco/Lubioscience) and added to 1 ml of Histopaque-1077, and then centrifuged at 400 x g for 30 minutes at room temperature, without the brake. Following centrifugation, the buffy coat was collected into 5 mL tubes and washed with PBS (400 x g for 10 minutes at 4°C). The cells were washed with FACS buffer stained with antibodies to detect surface expression of CD4 (GK1.5) and CD25 (PC61) (BD Biosciences, Franklin Lakes, NJ), whereas intracellular expression of FoxP3^+^ was stained using an anti-Mouse/Rat FoxP3 (FJK-16s) staining kit (eBiosciences, San Diego, CA). Intracellular staining for IL17A was performed using the intracellular fixation permeabilization kit and anti-mouse IL17 (TC11-18H10.1) specific antibodies from eBioscience™. Cells were also stained with antibodies to detect surface expression of CD45 (30-F11) and CD4 (clone GK1.5) to gate on the leukocyte population.

### T cell proliferation and cytokine assay

Mice were euthanized 10 days after immunization with PLP_91–110_ and draining lymph nodes were removed and challenged *in vitro* with PLP_91–110,_ as described previously (27). The results are presented as stimulation indices, which are counts per minute (cpm) of test sample/cpm of the control). For cytokine analysis, supernatants were collected from culture 48 h after peptide stimulation, and the concentration of cytokines (IL-17F) were measured by sandwich ELISA using pairs of relevant anti-cytokine monoclonal antibodies (Pharmingen, San Diego, CA).

### Isolation of CD4^+^CD25^+^ Treg, CD4^+^CD25^-^ T effector, and Dendritic cells

CD4^+^CD25^+^ T cells isolated from the spleens of DR3 and DR3.IL17A^-/-^ transgenic mice using the EasySep™ Mouse CD4^+^CD25^+^ regulatory T Cell Isolation Kit II (STEMCELL Technologies, Vancouver, Canada) according to the manufacturer’s protocols. CD4^+^CD25^-^ T effector cells were isolated from the Treg-depleted fraction of splenocytes using Anti-Mouse CD4 Magnetic Particles – DM (BD Biosciences, CA, USA). DCs were isolated from total splenocytes using CD11c microbeads according to the manufacturer’s protocols (BD Biosciences, CA, USA). The purity of specific cell populations was analyzed by flow cytometry; all populations used were >90% pure.

### *In vivo* depletion of CD4^+^CD25^+^ Treg cells

DR3 and DR3.IL17A^-/-^ transgenic mice received two intraperitoneal injections (200 µg/injection) of monoclonal anti-mouse CD25 (IL-2Ra) antibody (clone: PC-61.5.3; BioXcell) in 200 µl of PBS per dose on days-5 and −3 before induction of EAE (28). Controls received two intraperitoneal injections of 200 µl of PBS. CD4^+^CD25^+^FoxP3^+^ T cell depletion was confirmed by flow cytometric analysis of PBMCs on the day EAE was induced. For IL-17A depletion, DR3 transgenic mice were injected intraperitoneally with 100 ug/mouse anti-IL17A (Clone 17F3; BioXcell), IgG1 (Clone MOPC-21; BioXcell) or PBS. IL-17A was blocked in C57BL/6 mice by administering 100 ug/mouse anti-IL-17A, IgG1 or PBS by intraperitoneally injection every three days following induction of EAE, starting at day 5 through day 32 (e.g., days-24, −21, −18, −15, −12, −9, −6 and −3 pre EAE induction and post EAE induction (at day 5, 8, 11, 14, and 17). IL-17F or GM-CSF cytokine inhibition was performed *in vivo* in DR3.IL17A^-/-^ transgenic mice by intraperitoneally injection of 100µg/mouse anti-IL-17F (clone; MM17F-8F5; BioXcell) or 100µg/mouse of anti-GM-CSF (clone; MP1-22E9; BioXcell) post induction of EAE on days 5, 10, and 14. Control DR3.IL17A^-/-^ transgenic mice received 100µg/mouse IgG1 (Clone MOPC-21; BioXcell).

### Treg suppression assay

Spleens from 8-10-week-old DR3 and DR3.IL17A^-/-^ transgenic mice were harvested for CD4^+^ T cells by BD™ IMag Particles as per manufacturer instruction. T cell-depleted splenocytes were used as antigen presenting cells after UV irradiation. CD4^+^CD25^+^ T cells were isolated from splenocytes of DR3 and DR3.IL17A^-/-^ transgenic mice using EasySep™ Mouse CD4^+^CD25^+^ regulatory T cell isolation kit II (STEMCELL Technologies, Vancouver, Canada) according to the manufacturer’s protocols. Treg suppression assays were performed as described previously (29).

### Co-housing of DR3 and DR3.IL17A^-/-^ transgenic mice

For fecal transplantation, DR3.IL17A^-/-^ transgenic mice were recipients of fecal transplants from DR3 transgenic mice through oral gavage, and conversely, DR3 transgenic mice were recipients of fecal transplants from DR3.IL17A^-/-^ transgenic mice through oral gavage. Five days after oral gavage, DR3 transgenic mice were cohoused with DR3.IL17A^-/-^ transgenic mice. Fecal samples were collected before and after co-housing for shotgun metagenomic sequencing.

### Microbiota DNA isolation, sequencing, and analysis

Five female mice of each genotype, DR3 and DR3.IL17A^-/-^, were assessed for the composition of their gut microbiota before and after cohousing, resulting in a total of 20 fecal samples. DNA was extracted using PowerSoil DNA Isolation Kit (Qiagen) following the manufacturer’s protocol. Each metagenomic DNA was quantified and sequenced using an Illumina MiSeq resulting in more than 55 Gb of raw paired-end reads from 20 samples after shotgun metagenomic sequencing. The raw paired-end metagenomes were quality trimmed using Trim Galore version 0.5.0, followed by host read filtering by mapping to the mouse genome (GRCm39) using Bowtie2 (30), with default parameters resulting in 49.43 Gb of high-quality data. The quality trimmed and filtered forward and reverse reads were taxonomically characterized using MetaPhlAN 3.0 and mpa_v30_CHOCOPhlAN_201901 database with default parameter settings. The obtained relative percentages of the microbial taxa were compared between the groups using the Kruskal-Wallis test followed by Dunn’s multiple t-tests in GraphPad Prism. The filtered and trimmed sequences were further assembled using MegaHit (31). After assembly, Prokka was used to predict ORF based protein coding genes and assign functions to the predicted genes (32). BBMap aligner was used to produce quantifiable gene counts in the FPKM format (fragments per kilobase of gene per million) (https://jgi.doe.gov/data-and-tools/bbtools/). InterProScan was utilized for assignment for KEGG pathways and GO processes (https://www.ebi.ac.uk/interpro/search/sequence/). FPKM values of every protein/gene assigned to each of the KEGG pathway or GO process were derived from mapping the gene to the FPKM list of the genes identified in the previous step. Relative abundances of genes and pathways were used for downstream statistical analysis. Statistical significance between the groups were determined using non-parametric Wilcoxon signed rank test, and data was visualized using GraphPad Prism.

### In vivo depletion of IL-17A in C57BL/6 mice

C57BL/6 mice were placed in three groups that received antibody injections prior to EAE induction (day-24, −21, −18, −15, −12, −9, −6, and −3): the first group of mice received intraperitoneal injections (200 µg/injection) of monoclonal anti-mouse IL-17A (clone 17F3; BioXCell) in 200µl PBS per dose; the second group of mice received 200 µg/injection of isotype control (mouse IgG1) in 200µl PBS per dose; and the third group of mice received 200µl PBS to serve as vehicle control. Similarly, three groups of C57BL/6 mice were treated with either monoclonal anti-mouse IL-17A (200 µg/injection), isotype control mouse IgG1 (200 µg/injection), or PBS starting on day 5 post EAE induction, and continuing every third day until day 32. Mice were observed daily for disease symptoms and EAE was scored as per the scoring system described (33).

### Transcriptome sequencing and analysis

Tissue sections from the colon of both DR3.IL17A^-/-^ and DR3 mice before EAE induction were collected, chopped into small pieces, stored in RNAlater (Millipore Sigma, MO, USA), and frozen to −80^0^C until further use. RNA was extracted using the RNeasy Kit (Qiagen) following the manufacturer’s instructions. The quality and quantity of the RNA was evaluated using a NanoDrop and sequenced at Novogene Co. Ltd, CA. Sequences were quality controlled using default *fastqc* and paired FASTQ files were aligned to the mm10 genome build with the kallisto pseudoaligner (34) using 100 bootstraps. Estimated counts for transcripts were then aggregated into gene-level quantifications based on gene symbol. An average of 14.79 ±1.3 million transcripts were obtained for each sample. Further downstream processes were carried out in R 4.0.3 (35) using edgeR 3.32.1 (36) to identify differentially expressed genes, as described previously. Differentially expressed genes were fed into ShinyGO (37) to identify changes to genes involved in KEGG database using the mouse reference database.

### Quantification of PPAR Signaling Pathways Genes using qPCR

Tissue sections from the colon of 8-weeks old DR3.IL17A^-/-^ and DR3 mice were collected, and RNA was extracted using the RNeasy Kit (Qiagen) following the manufacturer’s instructions. The quality and quantity of the RNA was evaluated using a NanoDrop. To determine the expression of PPAR signaling pathway genes, namely *Adipoq*, *Plin1*, *Plin4*, and *Fabp4*, a two-step quantitative reverse transcriptase-mediated real-time PCR (qPCR) was performed. Equal aliquots of total RNA (2 µg) from colonic samples of DR3.IL17A^-/-^ (n = 5) and DR3 mice (n = 5) were reverse transcribed to cDNA using the High-Capacity iScript cDNA synthesis kit (Bio-Rad). qPCR reactions were then carried out in triplicate using 50 ng of cDNA and the Applied Biosystems Power SYBR Green PCR Master Mix (Applied Biosystems). Amplification data were collected using an Applied Biosystems Prism 7900 sequence detector and analyzed with Sequence Detection System software (Applied Biosystems). The qPCR primers for the genes were purchased from Integrated DNA Technologies, Inc. The primer sequences used were as follows: *Adipoq*: Fwd: AGATGGCACTCCTGGAGAGAAG, Rev: ACATAAGCGGCTTCTCCAGGCT *Plin1*: Fwd: GAGAAGGTGGTAGAGTTCCTCC, Rev: GTGTGTCGAGAAAGAGTGTTGGC *Plin4*: Fwd: GCACTAAGGACACGGTGACCAC, Rev: GACCACAGACTTGGTAGTGTCC *Fabp4*: Fwd: TGAAATCACCGCAGACGACAGG, Rev: GCTTGTCACCATCTCGTTTTCTC The data obtained from qPCR were normalized to the expression of the housekeeping gene GAPDH.

To assess the effect of *Prevotella copri* (*P. copri* DSM 18205) treatment on the expression of PPAR signaling pathway genes, DR3 mice were divided into two groups. One group of mice was treated with *P. copri*, while the other group was treated with BHI media (control). The *P. copri* group received 10^7^ CFUs of live *P. copri* DSM 18205 via oral gavage every other day for a total of seven doses. The control group received 200 µl of BHI media alone by oral gavage. Tissue sections from the colon of *P. copri*-treated and BHI media-treated DR3 mice were collected, and RNA extraction and qPCR analysis for the expression of PPAR signaling pathway genes (*Adipoq, Plin1, Plin4,* and *Fabp4*), Treg, and IL10 genes were performed as described above.

### Data and materials availability

All data needed to evaluate the conclusions in the manuscript are present in the manuscript and/or the Supplementary Materials. Shotgun metagenomic sequencing and RNA-seq data had been uploaded under bio-project PRJNA808406. No custom codes were used for the data analysis.

## Results

### IL-17A deficient HLA-class-DR3 transgenic mice develop milder EAE disease

In the HLA class-II transgenic model of EAE, mice lacking IFNγ develop severe brain-specific disease characterized by elevated levels of IL-17A (22). Thus, we hypothesized that in the absence of IL-17A (DR3.IL17A^-/-^), mice would be resistant to EAE. All transgenic mice developed normally with no obvious signs of pathology, and analysis of splenocytes confirmed that these mice lacked IL-17A-expressing CD4^+^ T cells (Fig. S1).

Interestingly, both DR3 and DR3.IL17A^-/-^ transgenic mice develop EAE disease with similar disease onset and incidence (Fig. 1A, 1B). However, DR3.IL17A^-/-^ transgenic mice developed milder disease over time and had a significantly lower cumulative clinical EAE and peak EAE score compared to DR3 mice (Fig. 1B, 1C, 1D). Histological analysis of the brain and spinal cord tissues demonstrated milder inflammation and demyelination in DR3.IL17A^-/-^ transgenic mice, consistent with decreased inflammatory cell infiltration in CNS (Fig. 1E, 1F, 1G, 1H). Thus, our data indicate that IL-17A is not required for the induction phase of the disease. However, the milder disease that occurred in the absence of IL-17A over time suggests that it may play a role in disease progression. To test this, we depleted IL-17A in DR3 transgenic mice by treating them with an anti-IL-17A antibody before induction of EAE (Fig. 2). We treated mice with the anti-IL-17A antibody on days −19, −14, −11, −8, and −4, before immunization with PLP_91-110_ for EAE induction (Fig. 2A). Compared to the control group DR3 transgenic mice pre-treated with anti-IL-17A still developed EAE however showed a milder disease course which was similar to EAE observed in DR3.IL17A^-/-^ transgenic mice (Fig. 2B, 2C). Thus, the development of milder disease in DR3 mice in the absence of IL-17A (genetic deficiency and IL-17A neutralization) suggests IL-17A is redundant for the induction phase of the disease but might be crucial for disease progression.

**Fig. 1:**
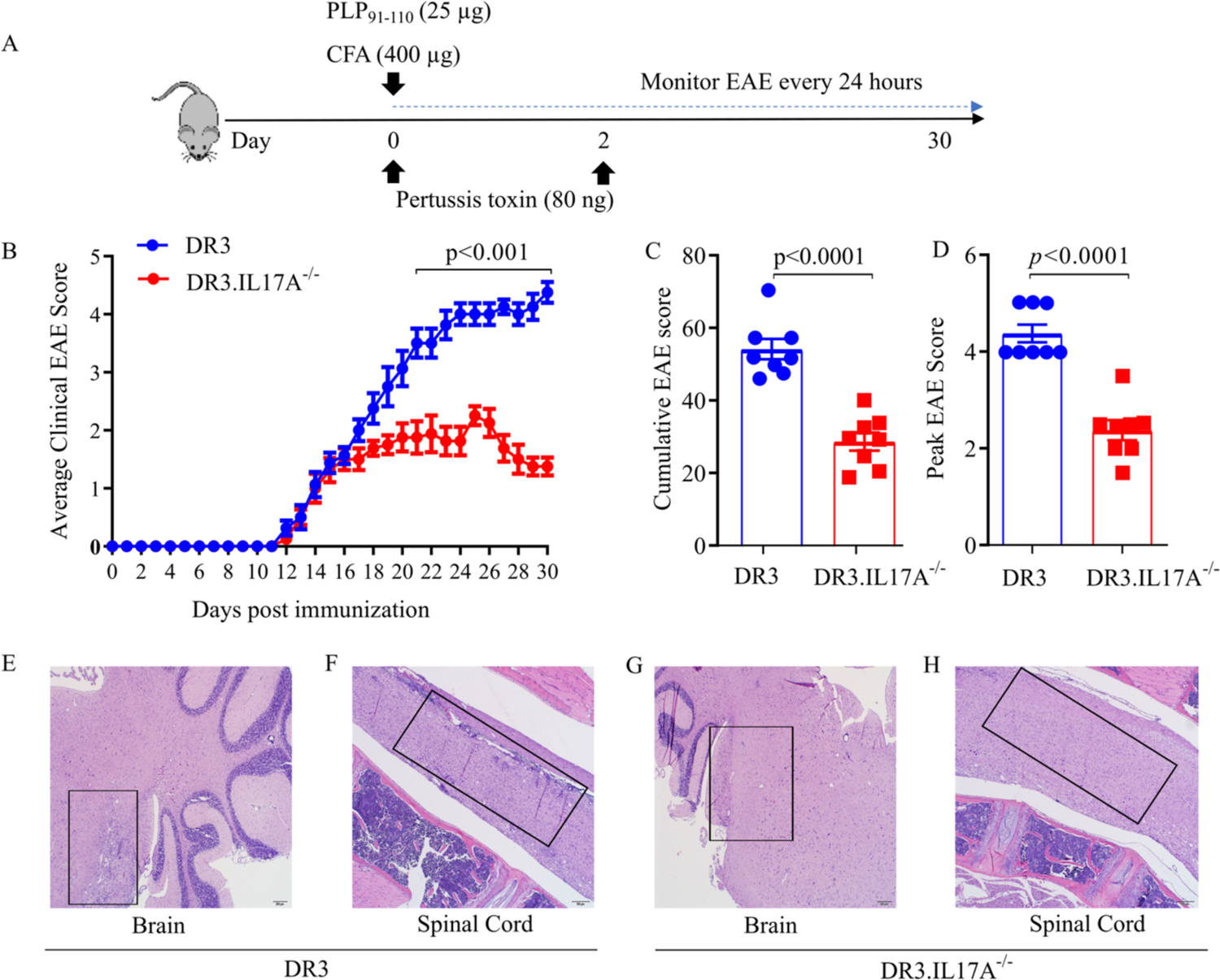
IL-17A deficiency ameliorates EAE in DR3 transgenic mice. (**A**) Schematic design of disease course in DR3 mice. DR3 (n= 8) and DR3.IL17A^-/-^ (n= 8) mice were immunized with PLP_91-110_ /CFA at day 0 and Pertussis toxin at days 0 and 2 to induce EAE. Mice were monitored for EAE disease progression before euthanizing on day 30 post-immunization. (**B**) Clinical EAE scores from mice in A, acquired over time. (**C**) Cumulative EAE scores from mice in A. (**D**) Peak EAE score from mice in A. (**E-H**) Representative histological images of tissues from the CNS from mice in A. Data are representative of 3 independent experiments with 4-5 mice per group. Bullet points (B) or bars (C, D) represent the mean and error bars represent the standard error of the mean for each experimental group. *p*-value was determined by multiple unpaired t-test (B), and unpaired t-test with Welch correction (C, D).

**Fig. 2:**
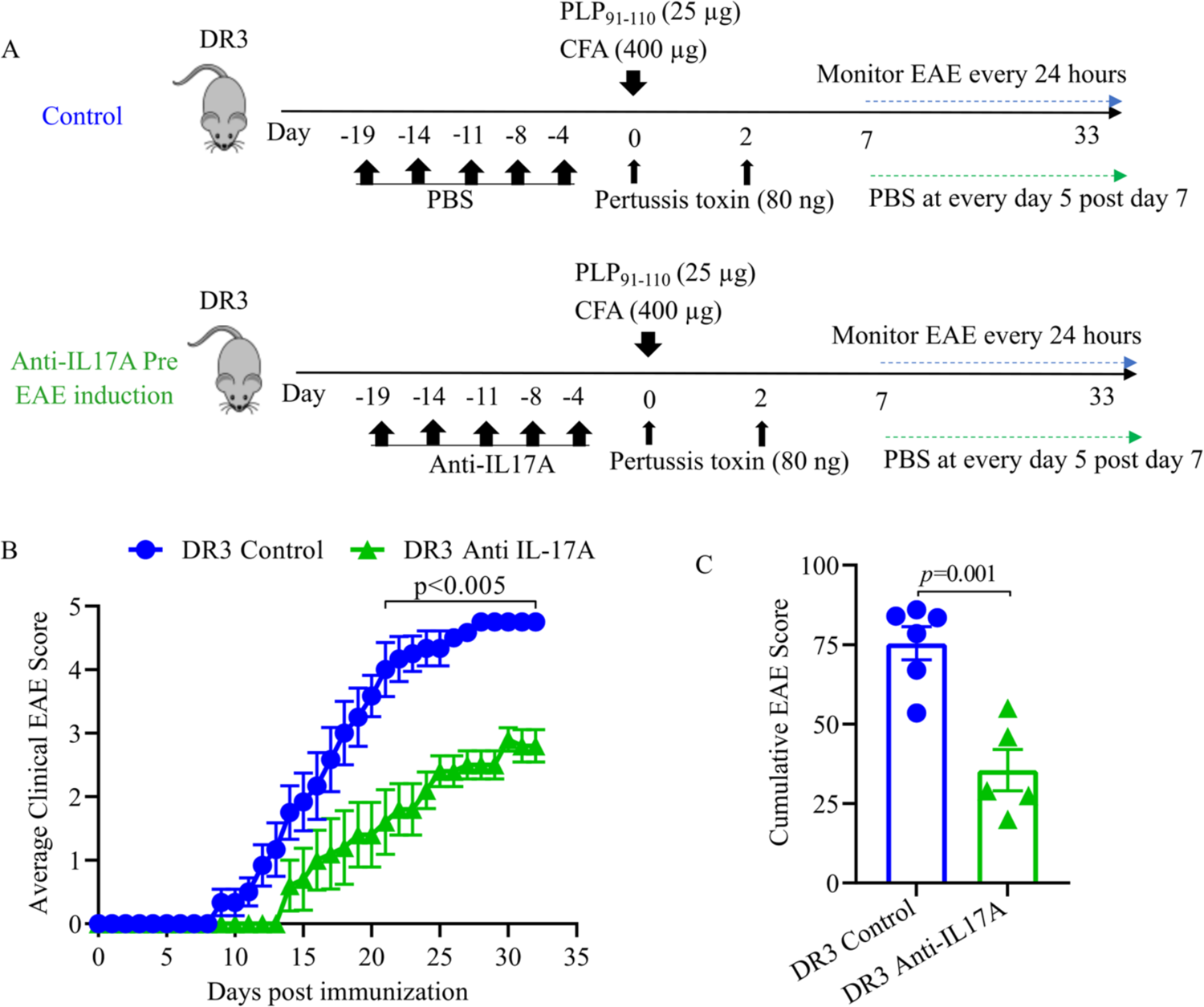
Anti-IL-17A treatment reduces the severity of EAE in DR3 mice. (**A**) Schematic design of anti-IL-17A treatment in mice induced with EAE. 8-10-week-old DR3 mice were treated with an anti-IL-17A monoclonal antibody before EAE induction with PLP_91-110_/CFA. (**B**) Average clinical EAE scores of mice treated as in A. Bullet points represent the mean and bars represent the standard error of the mean of each experimental group. (**C**) Average cumulative EAE scores of mice in A. Bars represent the mean and standard error of mean in each group. Data are from two independent experiments (n ≥4 mice per group). *p*-value determined by multiple unpaired t-test (B) and unpaired t-test with Welch correction (C).

To determine if the effect of IL-17A on EAE disease was universal or limited to the HLA class II transgenic mouse model, we assessed the effects of depleting IL-17A in wild-type C57BL/6J (B6) mice which require immunization with MOG_35-55_ for EAE (38). B6 mice treated with anti-IL-17A antibody after the onset of EAE (i.e., at days 5, 8, 11, 14, 17, 20, 23, 26, 29, and 32 post-disease induction) showed significantly lower average clinical scores over time compared to mice treated with PBS or IgG1a (Fig. S2A, S2B). Similar to DR3 transgenic mice, B6 mice that received pre-treatment (i.e., at days-24,−21,−18,−15,−12,−9,−6, and −3) and post-treatment (days 5, 8, 11, 14, and 17) of anti-IL-17A antibody still developed EAE but exhibited lower disease severity (Fig. S2C, S2D). These data indicated that the IL-17A is redundant for disease induction in both PLP-DR3-EAE and MOG-B6-EAE models. Neutralization of IL-17A exacerbated EAE progression in both HLA-DR3 and Wt B6 mice, suggesting a similar role of IL-17A in two models of MS disease.

Even though IL-17A is considered an important cytokine for the development of EAE (39), our results suggest that IL-17A is crucial for disease progression rather than disease induction. It is possible that IL-17F (shares 55% homology at the amino acid level with IL-17A) and other inflammatory cytokines such as GM-CSF or IFNγ might compensate for IL-17A to induce disease. We also found that IL-17A deficient mice have significantly higher levels of IL-17F compared to IL-17A sufficient DR3 transgenic mice (Fig. S3). However, neutralizing IL-17F did not affect disease incidence or severity (Fig. S4A, S4B, S4C). Next, we investigated whether GM-CSF was compensating for EAE induction in IL-17A deficient mice, as GM-CSF is shown to be a pathogenic cytokine in EAE (40). Although antigen specific CD4 T cells from DR3.IL17A^-/-^ transgenic mice did not show increased levels of GM-CSF but neutralization of the GM-CSF in DR3.IL17A^-/-^ resulted in an increased average disease score at a later point of the disease (Fig. S4A, S4B). However, the cumulative disease score was similar between isotype control and anti-GM-CSF treated mice (Fig. S4C). Thus, our data suggested that niether GM-CSF nor IL-17F can compensate for IL-17A in DR3.IL-17A^-/-^ transgenic mice for disease induction. Further, to determine the role of IFNγ in disease progression, we generated DR3 transgenic mice in which both IFNγ and IL-17A were deleted (DR3.IL17A^-/-^IFNγ^-/-^). Interestingly, these mice still developed diseases similar to those observed in DR3 transgenic mice (Fig. S5), suggesting that neither IL-17A nor IFNγ is required for the development of EAE. Taken together, our data indicate that neither IL-17A nor IL-17F nor GM-CSF nor IFNγ are required for the development of EAE disease. Thus, our data point toward an inflammatory mediator besides IL-17A, IL-17F, IFNγ or GM-CSF which is required for the induction of EAE disease.

### IL-17A deficiency enhances the proliferation of CD4^+^CD25^+^FOXP3^+^ Treg cells

The milder course of EAE disease observed in IL-17A deficient mice suggested the possibility of an immune-regulatory mechanism at play to dampen the chronic immune response. Studies have shown that CD4^+^CD25^+^Foxp3^+^ regulatory T cells can regulate autoimmune inflammation in the body and suppress IL-17A-secreting CD4+ Th17 cells (41, 42). Therefore, we analyzed the total number of CD4^+^CD25^+^Foxp3^+^ regulatory T cells in the peripheral blood of DR3 and DR3.IL17A^-/-^ mice. We observed that DR3.IL-17A^-/-^ mice had a significantly higher percentage (Fig. 3A, 3B) and number (Fig. 3C) of CD4^+^CD25^+^Foxp3^+^ regulatory T cells compared to DR3 mice. To determine whether the increased Treg levels were responsible for milder disease phenotype in DR3.IL17A^-/-^ mice, we treated these mice with an anti-CD25 blocking antibody to neutralize the Treg population as shown in Fig 3D and described previously (43). CD25 neutralization in DR3.IL17A^-/-^ mice resulted in severe disease phenotype similar to DR3 transgenic mice, indicating that Treg were involved in establishing milder disease phenotype in DR3.IL17A^-/-^ mice (Fig. 3E). Notably, CD4^+^CD25^+^ Treg that were isolated from the spleen of naïve DR3.IL17A^-/-^ transgenic mice had significantly higher suppressive ability against antigen specific CD4^+^CD25^-^ effector cells when compared to those of DR3 transgenic mice (Fig. 3F), further supporting the role of Treg in determining clinical EAE outcomes in DR3.IL17A^-/-^ transgenic mice. Collectively, our data indicate that deficiency of IL-17A results in a higher frequency and number of CD4^+^CD25^+^ Treg cells with a higher suppressive ability, which likely contributes to a milder disease phenotype in DR3.IL-17A^-/-^ transgenic mice.

**Fig. 3:**
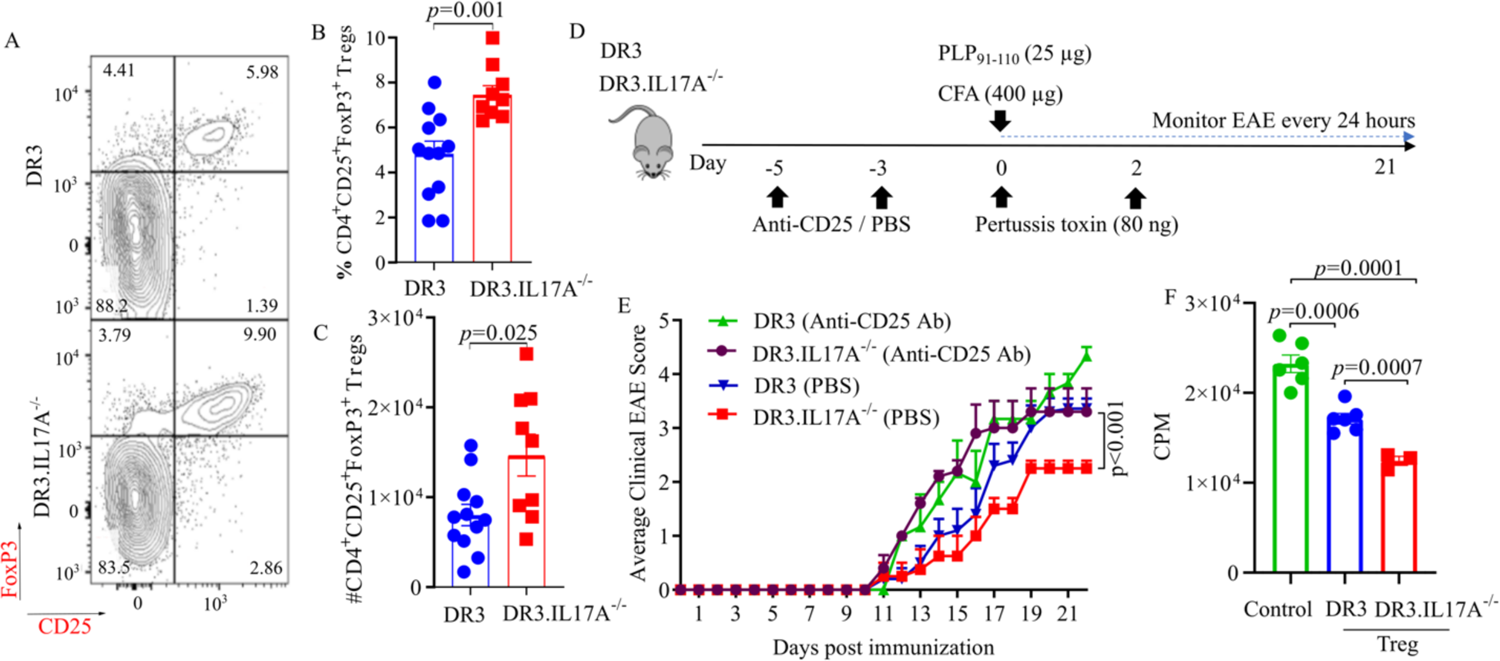
IL-17A deficiency in DR3 transgenic mice results in expansion of CD4^+^CD25^+^FOXP3^+^ Treg cells. (**A)** Representative flow cytometric plots of peripheral blood mononuclear cells (PBMCs) isolated from 8-10-week-old DR3 and DR3.IL17A^-/-^ transgenic mice. Plots were gated on live lymphocytes and singlets. (B) Percentage and (C) total number of CD4^+^CD25^+^FoxP3^+^ regulatory T cells from mice shown in A. (**D**) Schematic diagram of experimental workflow. DR3 and DR3.IL17A^-/-^ transgenic mice were treated with anti-CD25 to neutralize Treg prior to immunization with PLP_91-110_ /CFA to induce EAE. Arrowheads represent days in which mice received anti-CD25 treatment. (**E**) Average clinical EAE scores of DR3 and DR3.IL17A^-/-^ mice that were treated with anti-CD25 or PBS prior to induction of EAE. (**F**) Counts per minute (CPM) of DR3 CD4 effector cells and antigen presenting cells cultured with CD4^+^CD25^+^ Treg from the spleen of naïve DR3 or DR3.IL17A^-/-^ transgenic mice. The data represent two experiments performed at different time points. Bars represent the mean and standard error of the mean from two independent experiments. n =4-8 mice per group. *p*-value was determined by unpaired t-test with Welch correction (B, C, F) and two-way ANOVA for EAE clinical scores (E).

### IL-17A deficiency modulates the composition of the gut microbiome

The composition and function of the gut microbiota might impact the immune response. As the gut microbiome is known to modulate the Treg, first, we hypothesized that the absence of IL-17A modulated the composition of the gut microbiome, which then can induce the Treg population and lead to a milder disease phenotype in DR3.IL17A^-/-^ transgenic mice. Secondly, we hypothesize that the microbiota enriched in IL-17A deficient mice, when transferred to IL-17A sufficient mice, should alter the microbiota of DR3 transgenic mice to enhance Tregs in the gut.

To test our first hypothesis, we collected fecal samples from both DR3 and DR3.IL17A^-/-^ transgenic mice before cohousing and performed shotgun metagenomic sequencing to characterize the microbiome in these mice (Fig 4A). The microbial composition of DR3 transgenic mice clustered distinctly from that of DR3.IL17A^-/-^ mice (Fig. S6). Specifically, we observed a significant increase in the average relative abundance of *Prevotella* sp. MGM1, *Parabacteroides distasonis,* and *Bacteroides sartorii* in DR3.IL17A^-/-^ mice (Fig. 4B, 4C, 4D, 4E). We did observe that DR3.IL17A^-/-^ mice had a significant reduction in the population of *B. vulgatus*, *Lachnospiraceae* bacterium 10_1, *Lachnospiraceae* bacterium 28_4, *Lachnospiraceae* bacterium COE1, and *Lachnospiraceae* bacterium A2 (Fig. 4F, 4G, 4H, 4I, 4J). A complete list of bacteria showing trend but not reaching statistical significance have been provided in Figure S7.

**Fig. 4:**
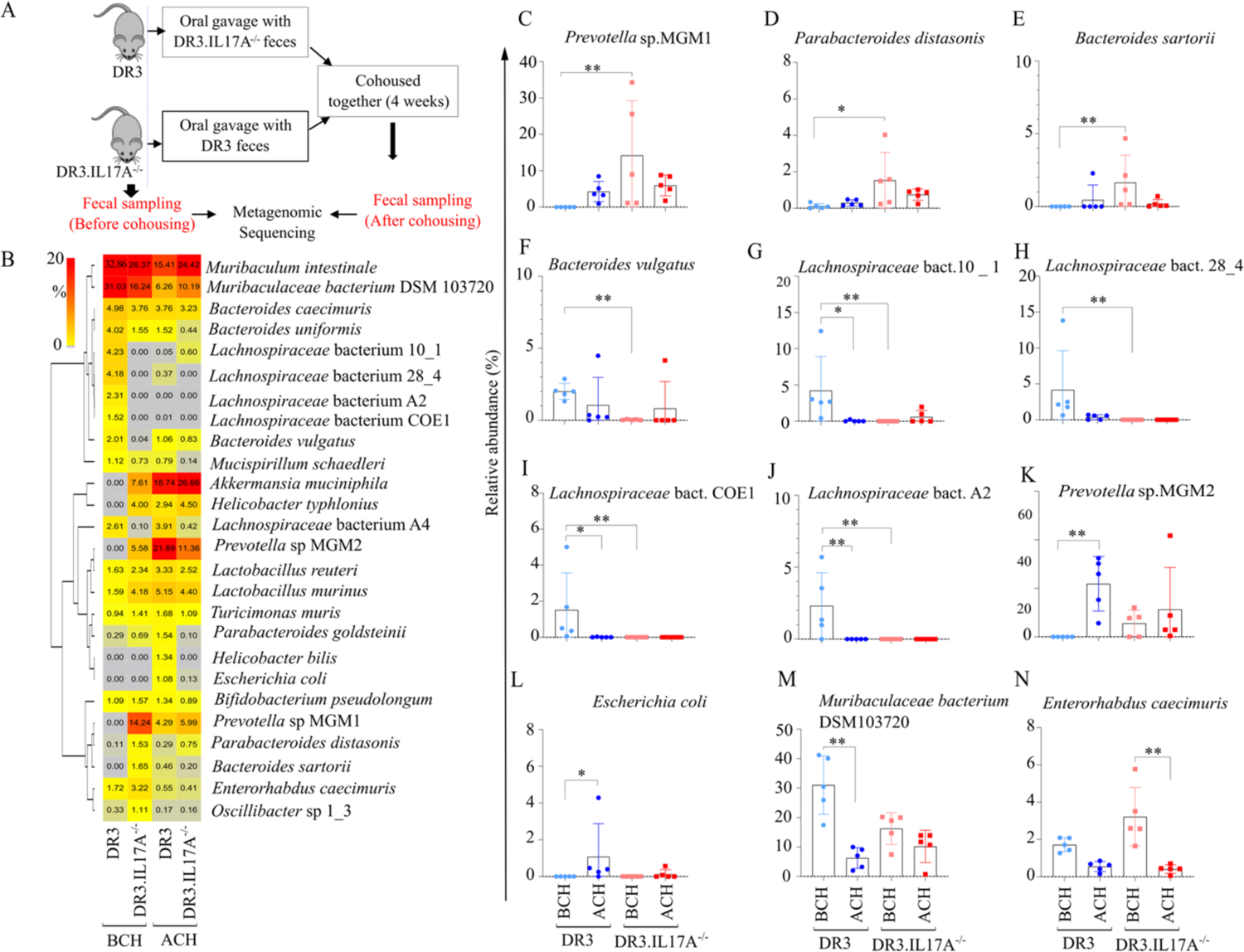
Taxonomic characterization of the gut microbiome in DR3 and DR3.IL17A^-/-^ mice. Fecal samples from DR3 and DR3.IL-17A^-/-^ transgenic mice before cohousing (BCH), or after that one time fecal transferred (orally gavaged) and cohoused (ACH) were obtained and subjected to shotgun metagenomic sequencing. (**A)** Schematic diagram representing the experimental flow. (B) Heatmap depicting bacterial species with > 1% average relative abundance in at least one group (n=5). (**C-N**). Statistically significantly abundant bacterial taxa that were statistically different in abundance in at least one of the comparisons (e.g., DR3-BCH to DR3.IL17A^-/-^-BCH, DR3-BCH to DR3-ACH, and DR3.IL17A^-/-^-BCH to DR3.IL17A^-/-^-ACH). Kruskal Wallis test followed by Dunn’s *posthoc* analysis was performed to identify significant differences between groups. “*” and “**” represent significance at *p*<0.05 and *p*<0.01 respectively.

For efficient microbiota transfer we first performed fecal transplantation through oral gavage followed by cohousing as described previously (44, 45). Microbiota analysis showed that the microbial composition in DR3 transgenic mice shifted to become more similar to that in DR3.IL17A^-/-^ transgenic mice (Fig. 4B, Fig. S6). Specifically, *Prevotella* sp. MGM2 (Fig. 4K) and *E. coli* (Fig. 4L) were present in DR3.IL17A^-/-^ mice were significantly increased in DR3 transgenic mice after cohousing with DR3.IL17A^-/-^ mice. *Prevotella* species were observed to be significantly higher in DR3.IL17A^-/-^ mice compared to DR3 transgenic mice prior to cohousing. Specifically, *Prevotella* sp. MGM2 had significantly higher relative abundance in DR3 transgenic mice after being cohoused with DR3.IL17A^-/-^ mice. Also, *A. muciniphila*, and *H. typhlonius* were absent in DR3 transgenic mice before they were cohoused but were increased but not significantly after cohousing with DR3.IL17A^-/-^ transgenic mice (Fig. 4B, Fig. S7). Similarly, the abundance of *Prevotella* sp. MGM1, *P. distasonis* and *B. sartorii* were increased, although not to significant levels, in DR3 transgenic mice after being cohoused with DR3.IL17A^-/-^ transgenic mice (Fig. 4C, 4D, 4E). In contrast, *Lachnospiraceae* bacterial species (Fig. 4G, 4I and 4J) and *Muribaculaceae* bacterium DSM 10370 (Fig. 4M) were significantly decreased in DR3 transgenic mice after being cohoused with DR3.IL17A^-/-^ transgenic mice. Thus, the cohousing of DR3 with DR3.IL17A^-/-^ transgenic mice transformed microbiota by transferring *Prevotella* species in DR3 mice.

### Cohousing DR3 mice with DR3.IL17A^-/-^ transgenic mice transfers a Treg-inducing phenotype

Next, we investigated whether that transferring the gut microbiome from DR3.IL17A^-/-^ mice to DR3 mice also affected Treg frequency and EAE severity. Therefore, we assessed the levels of Treg in the peripheral blood for both IL-17A deficient and IL-17A sufficient mice after fecal transplantation plus cohousing (Fig. 5A). We observed that CD4^+^CD25^+^FoxP3^+^ T cell populations were significantly increased in DR3 transgenic mice after cohousing with DR3.IL17A^-/-^ mice (Fig. 5B). In contrast, Treg populations were similar in DR3.IL17A^-/-^ mice cohoused with DR3 mice (Fig. 5C). These data suggest that microbes transferred from DR3.IL17A^-/-^ transgenic mice to DR3 transgenic mice such as *Prevotella* species were crucial to enhance the Treg population.

**Fig. 5:**
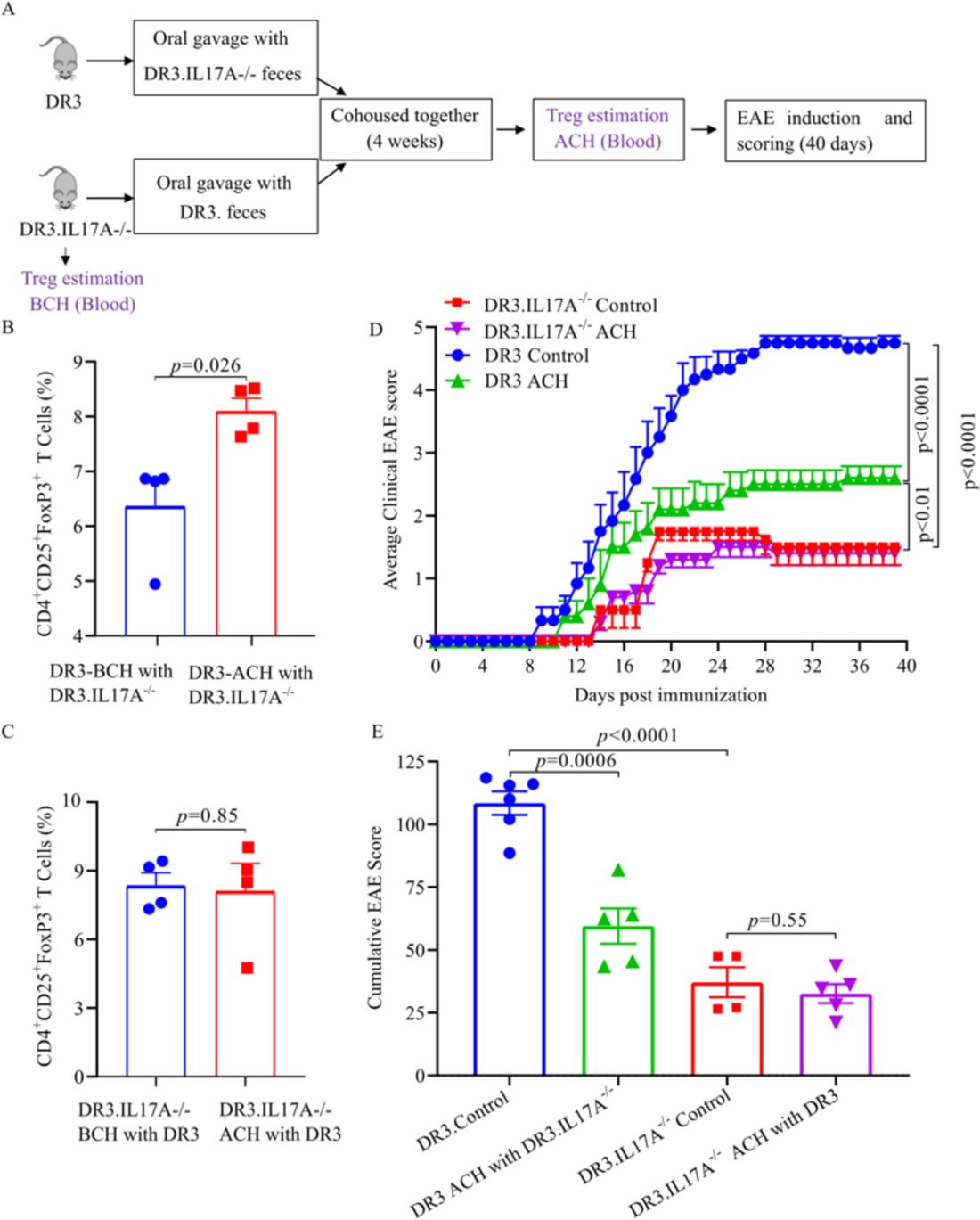
Treg-inducing bacteria from DR3.IL17A^-/-^ mice reduce disease severity when transferred to DR3 mice. **(A)** Schematic of experimental design showing experimental outline. 8-10-week-old female DR3 transgenic mice were cohoused for four weeks with DR3.IL17A^-/-^ transgenic mice after one-time fecal transferred (orally gavaged) from each other mice strains. Peripheral blood mononuclear cells (PBMCs) were isolated from blood that was collected by retro-orbital bleeding, and regulatory T cells were assessed by flow cytometric analysis. Percentages of CD4^+^CD25^+^FoxP3^+^ T cell populations in (**B**) DR3 transgenic mice before (BCH) and after (ACH) being cohoused with DR3.IL17A^-/-^ transgenic mice and in (**C**) DR3.IL17A^-/-^ transgenic mice before (BCH) and after (ACH) being cohoused with DR3 transgenic mice. (**D**) Average clinical EAE scores of DR3 transgenic mice (non-cohoused control and after cohoused with DR3.IL17A^-/-^ transgenic mice) and DR3.IL17A^-/-^ transgenic mice (non-cohoused control and after cohoused with DR3 transgenic mice) that were immunized with PLP_91-110_/CFA and followed for 30 days. (**E**) Average cumulative EAE scores of mice in C. Data represent the mean and standard error of the mean for each experimental group from two independent experiments: n ≥4 mice per group. *p*-value was determined by unpaired t-test with Welch correction (A, B, D), two-way ANOVA for clinical EAE scores (C).

Next, we examined whether cohousing also affected the outcome of the EAE disease phenotype. DR3 and DR3.IL17A^-/-^ transgenic mice were cohoused for four weeks and then housed separately while EAE was induced, and the course of the disease was assessed. When housed separately, DR3 mice showed more severe disease than DR3.IL17A^-/-^ mice as expected (Fig. 5D, 5E). In contrast, DR3 mice cohoused with DR3.IL17A^-/-^ mice showed milder disease compared to DR3 control mice that were never cohoused (Fig 5D, 5E). However, the disease in DR3.IL-17^-/-^ transgenic mice were similar pre and post-cohousing with DR3 transgenic mice. Thus, our results confirmed that the lack of IL-17A alters the composition of the gut microbiota to allow the expansion of bacteria such as *Prevotella* species, leading to the expansion of the Treg population and ultimately, to milder EAE disease phenotype.

### Short-chain fatty acid metabolic pathways are transferred from DR3.IL17A^-/-^ transgenic mice to DR3 transgenic mice after cohousing

Next, to identify the mechanism through which IL-17A-influenced gut microbiota can modulate Treg, we compared bacterial metabolic pathways enriched in DR3.IL-17A^-/-^ mice. We first assessed short-chain fatty acids (SCFAs), as these have been previously reported to affect the differentiation of T cells (46) and ameliorate EAE severity (47). The metabolic pathways associated with pyruvate to propanoate (P108-PWY) and pyruvate fermentation to acetate and lactate II (PWY-5100) trended higher, although not to significant levels, in DR3.IL17A^-/-^ transgenic mice compared to DR3 transgenic mice before cohousing. P108-PWY was low in abundance in both DR3 and DR3.IL17A^-/-^ transgenic mice before cohousing but was significantly higher in both strains after cohousing (Fig. S8A). Likewise, pyruvate fermentation to acetate and lactate II was lower in DR3 transgenic mice but became significantly enriched after mice were cohoused with DR3.IL17A^-/-^ transgenic mice (Fig. S8B). The transfer of SCFA pathways from DR3.IL17A^-/-^ transgenic mice to DR3 transgenic mice, but not vice versa, indicates that the DR3.IL17A^-/-^ transgenic mice harbor microbes that are responsible for SCFAs production and these microbes can be transferred to DR3 transgenic mice through co-housing. Thus, our data suggests the potential role of microbiota mediated SCFAs production in inducing Treg to produce milder disease in the absence of IL-17A.

### Loss of IL-17A enriches PPAR signaling genes in the colon of DR3 transgenic mice

The majority of immune cells reside in the gut to protect its large surface area, and gut bacteria play a vital role in generating these immune cells, especially Treg and Th17 cells. The importance of gut microbiota in generation of Treg is highlighted by studies performed in germ-free/antibiotic treated mice, which showed significantly reduced Treg population (48, 49). Therefore, to identify whether the host genes involved in modulating the disease in IL-17A deficient mice, we performed transcriptome analysis in the colonic tissues from naive DR3 and DR3.IL17A^-/-^ mice (Fig. 6A). We observed that the colonic tissues from mice of the different genotypes (e.g., DR3 and DR3.IL17A^-/-^) were distinctly separated (Fig. 6B). Upon differential analysis, we identified 123 genes that were differentially expressed in the colon between the two groups (Fig. 6C, Table S1).

**Fig. 6:**
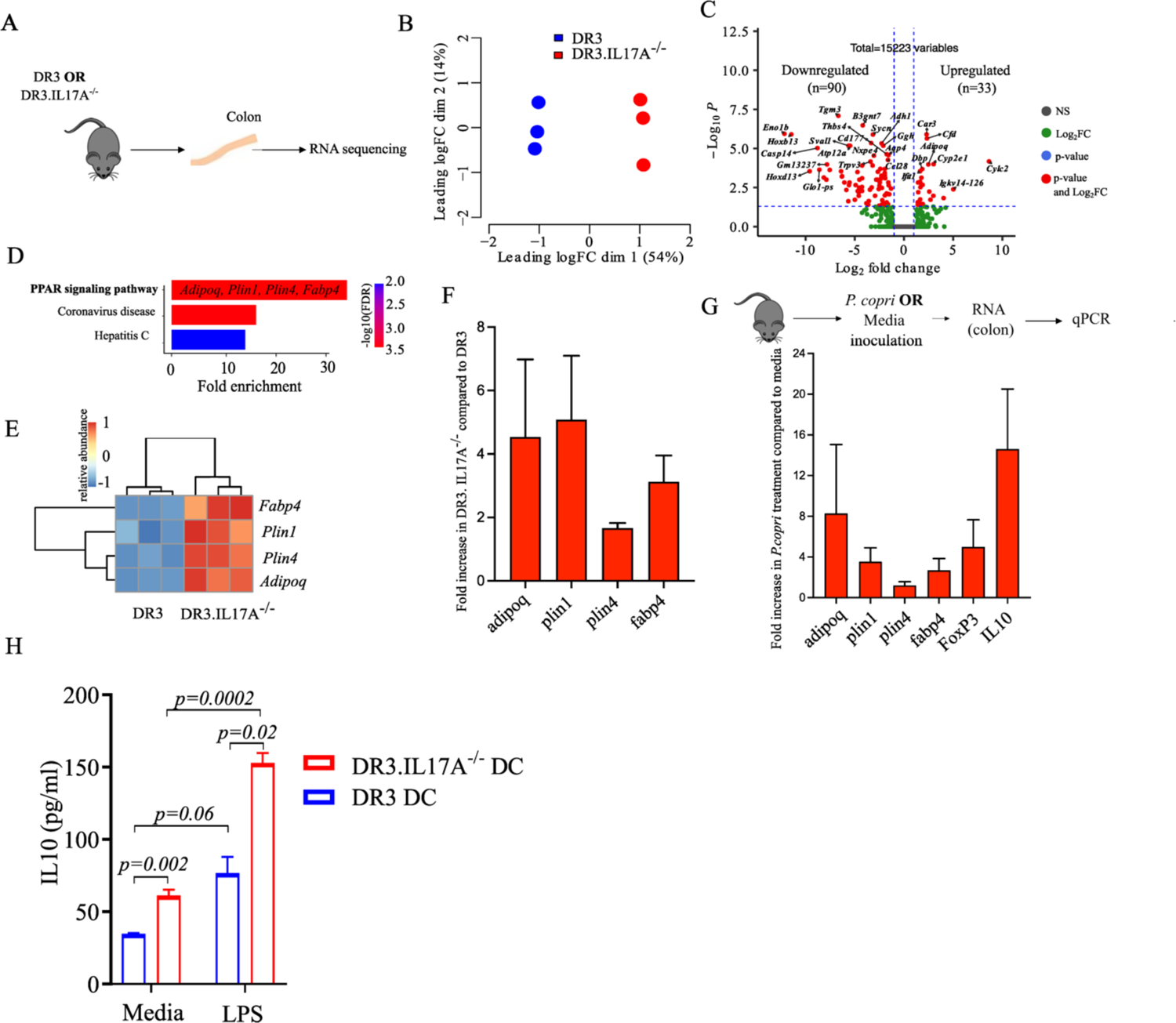
DR3 and DR3.IL17A^-/-^ transgenic mice exhibit distinct anti-inflammatory gene expression patterns in the colon. Colonic tissue was harvested from DR3 (n=3) or DR3.IL-17A^-/-^ transgenic mice (n=3) and subjected to RNA-seq analysis. (**A**) Schematic diagram of workflow. B) Multi-dimensional scaling plot showing the separation of DR3 and DR3.IL17A^-/-^ colonic samples. Each letter represents an RNA-Seq sample. Sample groups are indicated by different colors, as indicated in the legend. (**C**) Volcano plot displaying the pattern of gene expression of DR3.IL17A^-/-^ mice relative to DR3 mice. Genes that are significantly differentially expressed, with a Log_2_ fold change of >1 and an FDR-corrected p-value <0.05, are highlighted in red, blue lines represents the boundary for Log_2_ fold change and the adjusted p-value cutoff. D) Pathways enriched for 33 upregulated genes in KEGG database identified using ShinyGO webserver (37). E) Heatmap of enriched genes in PPAR signaling pathway. F) Ratio of average fold change expression of genes using qPCR related to PPAR signaling pathway in DR3.IL-17A^-/-^ transgenic mice to DR3 transgenic mice. G) Effect of *Prevotella copri* DSM 18205 supplementation in the colon of DR3 mice. Mice (n = 5 in each group) were treated with either *P. copri* DSM 18205 (10^7^ CFU) or media (control) for every alternate day for two weeks. After two weeks, mice were euthanized, and RNA was isolated from the colonic tissue. Genes related to PPAR pathways along with FoxP3 and IL-10 were quantified using qPCR. Ratio of average fold change expression of genes related to PPAR signaling pathway along with FOXP3 and IL10 in *P. copri* treated to media treated mice. (H) Quantification of IL-10 cytokine production in naïve dendritic cells from DR3 and DR3.IL17A^-/-^ transgenic mice following stimulation with LPS. Splenocytes were harvested from naïve 8-10-week-old HLA-DR3 and DR3.IL17A^-/-^ transgenic mice and CD11c+ dendritic cells were isolated using BD™ IMag Particles. Purified CD11c+ cells were activated with LPS for 24 hours and IL-10 was measured by ELISA. Media without LPS was used as control. Bars represent the mean and standard error of the mean from each experimental group; n ≥3 mice per group. p-value was determined by pairwise Mann-Whitney test.

Thirty-three out of 123 differentially expressed genes were upregulated in DR3.IL17^-/-^ mice relative to DR3 transgenic mice. PPAR signaling pathway was enriched upon KEGG database search consisting of *Adipoq, Plin1, Plin4,* and *Fabp4* genes (Figure 6D, 6E), which was sufficient to distinguish between the DR3 and DR3.IL17^-/-^ mice. These genes were further validated using qPCR where *Plin1, Plin4* and *Fabp4* and *Adipoq* were enriched with an average 5.07, 1.66, 3.12 folds in DR3.IL17A^-/-^ transgenic mice compared to DR3 (Fig. 6F). PPAR signaling mainly involves lipid synthesis, degradation, storage, transport, binding, oxidation, and cholesterol metabolism. Interestingly, activation of PPAR pathways is shown to enhance Treg responses (50), and inhibit the inflammatory response, producing significant anti-atherosclerosis (51) and Sjogren Syndrome-like Dacryoadenitis (52) by modulating Th1/Th17 response. Especially, *Plin1* and *Plin4* are involved with lipid storage, *Fabp4* with lipid binding, and *Adipoq* with lipid degradation and previous study has demonstrated that lipid metabolism is closely associated with improvements of mucosal barrier functions (53). Taken together, the upregulated genes in DR3.IL17A^-/-^ mice related to the PPAR signaling pathway suggest enrichment of Tregs exert an anti-inflammatory effect in the colon. On the other hand, for the downregulated genes, pathways belonging to alpha-linolenic acid and linoleic acid metabolism were enriched (Fig. S9A), and three genes related to these pathways were sufficient to distinguish DR3 from DR3.IL17A^-/-^ transgenic mice (Fig. S9B).

Our microbiome and transcriptome data both indicated that there is enhanced Treg response in the gut of DR3.IL17A^-/-^ transgenic mice through transfer of *Prevotella* species. We have previously shown that *Prevotella* species such as *P. histicola* increased the frequency and number of CD4+FoxP3+ regulatory T cells in gut (54). Thus, we hypothesize that *Prevotella* species will induce an anti-inflammatory effect in the colon by enhancing genes related to the PPAR signaling pathway and finally affecting the Treg population. Hence, DR3 mice were either gavaged with *P. copri* DSM 18205 (10^7^CFU) or media control every alternate day for two weeks. Upon quantifying the genes with qPCR, we observed that *Adipoq, Plin1* and *Fabp4* and *Plin4* genes were 8.26, 3.5, 1.2 and 2.67 average folds enriched in *P. copri* gavaged mice (Fig. 6G) suggesting that *P. copri* may have induced the PPAR signaling pathway. Interestingly, FoxP3 and IL-10 were 4.98 and 14.6 average folds enriched in *P. copri* administered mice compared to control (Fig. 6G), suggesting that *P. copri* treatment exerts anti-inflammatory effects in the colon by enhancing Tregs and IL-10.

### Dendritic cells from DR3.IL17A^-/-^ transgenic mice produce higher levels of IL-10

We observed the transfer of *Prevotella* species and enrichment of Tregs in DR3 after cohousing with DR3.IL17A^-/-^. Thus, we asked if the antigen-presenting cells, such as dendritic cells (DCs) in DR3 and DR3.IL17A^-/-^ are primed by the gut microbiota to become tolerogenic. Therefore, we analyzed the production of the anti-inflammatory cytokine IL-10 from LPS stimulated DCs from naïve DR3 and DR3.IL17A^-/-^ mice. Interestingly, CD11c^+^ DCs from naïve DR3.IL17A^-/-^ mice produced significantly higher levels of IL-10 compared to DR3 mice after stimulation with LPS (Fig. 6H). Thus, CD11c+ DCs from DR3.IL17A^-/-^ mice displayed a tolerogenic phenotype, marked by increased IL-10 production and trended reduced antigen presentation, potentially contributing to the milder disease course observed in these mice.

## Discussion

IL-17A has been shown to play an important role in the pathobiology of autoimmune diseases, including MS (4, 55, 56); however, the precise mechanism remains poorly understood. Here, we demonstrate the crucial role of IL-17A in regulating the progression of EAE, an animal model of MS. Our data showing milder disease but similar disease incidence and severity in DR3.IL17A^-/-^ mice indicate an important role of IL-17A in regulating disease severity through modulation of Treg populations and Treg-promoting gut bacteria like *Prevotella* species. The importance of Treg-promoting gut bacteria was further validated by gut microbiota transfer utilizing cohousing experiments. Specifically, co-housing IL-17A-sufficient DR3 transgenic mice with IL-17A-deficient DR3.IL17A^-/-^ transgenic mice altered gut microbiota composition and functions, enrichment in the Treg subset, and a milder disease phenotype similar to that observed in DR3.IL17A^-/-^ transgenic mice. Furthermore, an analysis of host gene expression in the colon of DR3.IL17A^-/-^ transgenic mice compared to DR3 transgenic mice uncovered a crosstalk between *Prevotella* species, the PPAR pathway, and Tregs in the gut. Thus, our results demonstrate an intricate interplay between the host and gut microbes in relation to IL-17A, which ultimately impacts the pathogenicity of central nervous system inflammatory diseases, such as MS.

The development of milder disease in DR3.IL17A^-/-^ transgenic mice suggest induction of regulatory pathways in the absence of IL-17A. In support of this, we observed an increased CD4^+^CD25^+^FoxP3^+^ Treg population in DR3.IL17A^-/-^ transgenic mice. Further, the disease was restored after Treg depletion using an anti-CD25 (PC61) antibody, confirming an important role for CD4^+^CD25^+^ Treg subset in disease regulation. Interestingly, similar observations have also been reported in GM-CSF deficient mice, which are resistant to EAE development, but treatment with anti-CD25 antibody (PC61) resulted in the development of severe disease (57). Thus, these data suggest that pathogenic cytokines such as IL-17A and GM-CSF might influence EAE disease progression by downregulating regulatory cells such as CD4^+^CD25^+^ Treg cells. Development of disease in mice lacking IL-17A or receiving CD4^+^ T cells from *Csf2*-/- (mice lacking GM-CSF) (57) supports our data that neither IL-17A nor GM-CSF are essential for disease development in the EAE model.

Interestingly, we observed that IL-17A significantly affects disease progression without affecting the onset of disease. This is consistent with other studies such as Haak et al., who showed that either loss of IL-17A or overexpression of IL-17A in CNS has a major impact on disease development (58). It is possible that other Th17 cytokines (IL-17F and GM-CSF) or even Th1 cytokine (IFNγ) can compensate for IL-17A loss for disease development. However, in our study, mice that were treated with blocking antibodies against IL-17F or GM-CSF still developed the disease. IL-17F is a member of the IL-17 family of cytokines, and several studies have investigated its role in EAE. However, the majority of studies have concluded that IL-17F is either redundant or does not play a role in EAE model (58, 59). IL-17F deficient mice treated with anti-IL-17A blocking antibodies were susceptible to EAE (58). The role of GM-CSF in EAE and MS is still a matter of debate, with some studies suggesting it is essential for EAE disease as GM-CSF deficient mice are resistant to EAE development (60) whereas others have shown that GM-CSF is not essential for EAE development (40, 61). Interestingly, Duncker et al., observed that although GM-CSF is required for disease progression, it was dispensable for disease induction phase (40). Development of severe disease in DR3 mice lacking both IL-17A and IFNγ ruled out the role of IFNγ in disease induction in the absence of IL-17A. Thus, our data suggest that IL-17A, together with IL-17F, GM-CSF, or IFNγ, is dispensable for the disease induction phase as DR3.IL-17A^-/-^ transgenic mice still develop the disease, although in a milder course. Future studies will be necessary to determine the mechanisms driving disease in the absence of IL-17A.

MS is driven by both host and environmental factors including gut microbiota, thus, both gut microbiota and host genes can contribute to increased Treg population in DR3.IL17A^-/-^ transgenic mice. In recent years, gut microbiota has gained prominence as a significant contributor to the induction and regulation of both Treg and Th17 cell subsets (49). Yet, our understanding of how IL-17A affects the gut microbiome remains limited, and there have been suggestions that IL-17A/F may function as modulators of intestinal microbiome and homeostasis (55). Indeed, in IL-17A deficient DR3 transgenic mice, we observed an increase in abundance of gut bacteria such as *Prevotella* species, *Parabacteroides distasonis,* and *Bacteroides species,* which are recognized for their capacity to promote Treg population (33, 54, 62, 63). Specifically, *Prevotella* sps. MGM2 successfully transferred to the IL-17A sufficient mice from IL-17A deficient mice in high numbers. Simultaneously, after cohousing IL-17A-deficient mice with IL-17A-sufficient mice, none of the microbial species’ abundances displayed significant alterations in IL-17A deficient mice. After fecal transplantation or cohousing, the gut bacteria in DR3 mice became more similar to that of DR3.IL17A^-/-^ mice. This suggests that the gut microbiota of DR3.IL17A^-/-^ mice is more resistant to change, unlike the gut bacteria in DR3 mice. Ecologically, the degree of resistance of the microbiota against incoming microbial species is crucial in determining health and disease (64). The reduced stability and loss of resistance in the gut microbiota of DR3 transgenic mice was reflected in increased disease severity, which was reduced upon cohousing with DR3.IL17A^-/-^ transgenic mice. This suggests that the gut microbiota in DR3 transgenic mice underwent reconfiguration to restore protective functions such as SCFAs production. SCFAs pathways producing acetate, propionate, and lactate were present in DR3.IL17A^-/-^ transgenic mice but not in DR3 transgenic mice. SCFAs produced by gut bacteria are involved in enrichment/proliferation of IL-10 producing Tregs and are crucial for maintaining gut homeostasis (65). In fact, the induction of Treg in IL-17A deficient mice was concurrent with the increased production of IL-10 from splenic DCs from DR3.IL17A^-/-^ transgenic mice. It suggests that the tolerogenic DCs are induced in the absence of IL-17A, and DC-derived IL-10 is essential to ameliorate EAE severity (66). The reduced trends of antigen presentation capacity of DCs from DR3.IL17A^-/-^ transgenic mice further support the role of tolerogenic DCs in mice lacking IL-17A. Even, SCFAs such as propionate and butyrate have been demonstrated as potent inducers of tolerogenic DCs (67). Thus, SCFAs produced by the gut microbiota like *Prevotella* species in DR3.IL17A-/-transgenic mice can induce tolerogenic DCs in IL-17A deficient mice. However, details of how the gut microbiota affects DC proliferation and IL-10 production need to be investigated further.

On the host side, IL-17A-deficient mice, characterized by an increased Treg population and altered gut microbiota composition, exhibited increased expression of PPAR signaling genes in our transcriptome analysis. Genes belonging to PPAR signaling categories are indicated as important mediators of anti-inflammatory responses (53, 68). Although a prior study involving both IL-17A and IL-17F found no impact on Treg population (55), our findings reveal markedly elevated levels of CD4^+^CD25^+^FoxP3^+^ T cell populations in the DR3.IL17A^-/-^ transgenic mice compared to DR3 transgenic mice suggesting a likely role of gut microbiome mediated Treg stimulation and anti-inflammatory effect in the colon. *Prevotella* species were enriched in DR3.IL17A^-/-^ transgenic mice were later transferred through cohousing to DR3 transgenic mice, amplified Tregs in DR3 transgenic mice; we investigated if *Prevotella* species can, in fact, induce PPAR genes and Tregs. We observed that *P. copri* DSM 18205 treatment to DR3 transgenic mice enriched colonic PPAR pathways genes. Interestingly, along with enrichment of PPAR associated genes, *P. copri* treatment to DR3 mice enriched FoxP3 and IL-10 gene expression in the colonic tissue. These findings suggesting that *P. copri* treatment exerts anti-inflammatory effects in the colon by enhancing the PPAR signaling pathway linked to Tregs and IL-10 induction. We have also previously shown that introducing *Prevotella* species can alleviate EAE severity through changes in the gut microbiota that ultimately promote Treg development (33, 54). Taken together, our study demonstrates gut microbiota-dependent internal mechanisms to maintain intestinal homeostasis and Treg function, and perturbation of the same can promote inflammation and CNS autoimmunity. Suppression of disease in IL-17A sufficient mice on cohousing with IL-17A deficient mice strongly suggests that Treg-promoting gut microbiota is dominant over disease-promoting bacteria and thus has strong clinical translation potential. However, the microbial community network and the interplay between the microbes in the community to exacerbate or protect from EAE warrants deeper exploration.

In summary, our study highlights an important pathway through which IL-17A regulates inflammation, specifically by modulating the gut microbiota to suppress Treg expansion. Our study illuminates a novel mechanism by which immune mediators such as cytokines impact the gut microbiota to alter immune cell function and, ultimately, disease outcomes. Our results have strong clinical relevance as modulation of the gut microbiota are being considered as potential therapeutics to treat diseases ranging from inflammatory to neurological diseases. As MS is an inflammatory disease characterized by gut dysbiosis, combination therapies targeted to reduce inflammation as well as correct gut dysbiosis might result in better therapeutic outcome than either one alone. Although IL-17A had been linked with the pathobiology of MS, two anti-IL-17A clinical trials in MS (NCT01874340 and NCT01433250) were terminated early. In that regard, while anti-IL-17A therapy alone was not as effective as current MS drugs, a combination of anti-IL-17A and microbiota modulation might provide better therapeutic efficacy. Additionally, Treg induction is also being tried as a potential treatment option in MS (69). Our cohousing experiment showing the ability of gut microbiota to colonize, induce Tregs, and suppress disease in susceptible mice suggests that gut microbiota-based induction of host immunity might have an added advantage as it will not only induce Tregs but can correct gut dysbiosis, too. Future studies testing gut commensal with Treg-inducing capabilities in combination with anti-IL-17A are warranted to test whether a combination of anti-inflammatory plus gut commensal modulation might provide added benefits.

## Acknowledgements

We thank the Waldschmidt, Legge, Jabbari, and Lieberman laboratories for helpful discussion. We thank Jian Zhang and John Harty for reading the manuscript and providing their valuable input. We also thank the Comparative Pathology Laboratory at the University of Iowa, Department of Pathology, for their time and efforts.

## Author Contributions

Conceptualization: AKM, SKS, NJK

Formal analysis: SKS, SG, NB, KNGC, SMG

Methodology: AKM, SKS, NB, NJK

Investigation: AKM, SKS, SNJ, SG, AGR, TT, NB, PL, AGR, TT, KNGC

Visualization: AKM, SKS, SNJ, SG, TT, NB, KNGC, SMG

Funding acquisition: AKM

Project administration: AKM, SKS

Supervision: AKM, NJK

Writing – original draft: AKM, SKS, SG,

Writing – review & editing: AKM, SKS, SG, SMG, NJK

## Supplementary Figures

**Fig. S1:**
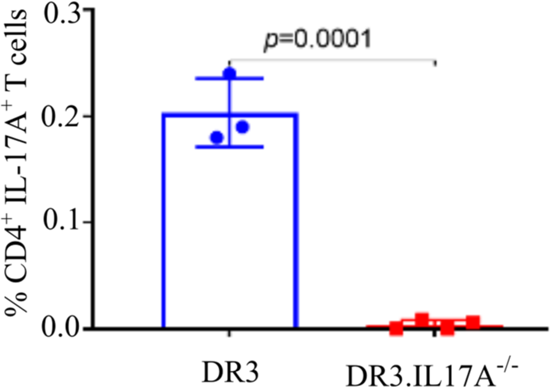
DR3.IL17A^-/-^ mice lack IL-17A-expressing CD4+ T cells in the spleen. 8-10-week-old DR3 and DR3.IL17A^-/-^ transgenic mice were euthanized and splenocytes were harvested for analysis. Quantification of the frequency of CD4^+^IL-17A^+^ T cells from mice in A. DR3: n=3; DR3.IL17A^-/-^: n = 4. *p*-value was determined using Mann-Whitney test between the experimental groups. Symbols represent individual mice, bars represent mean±SEM.

**Fig. S2:**
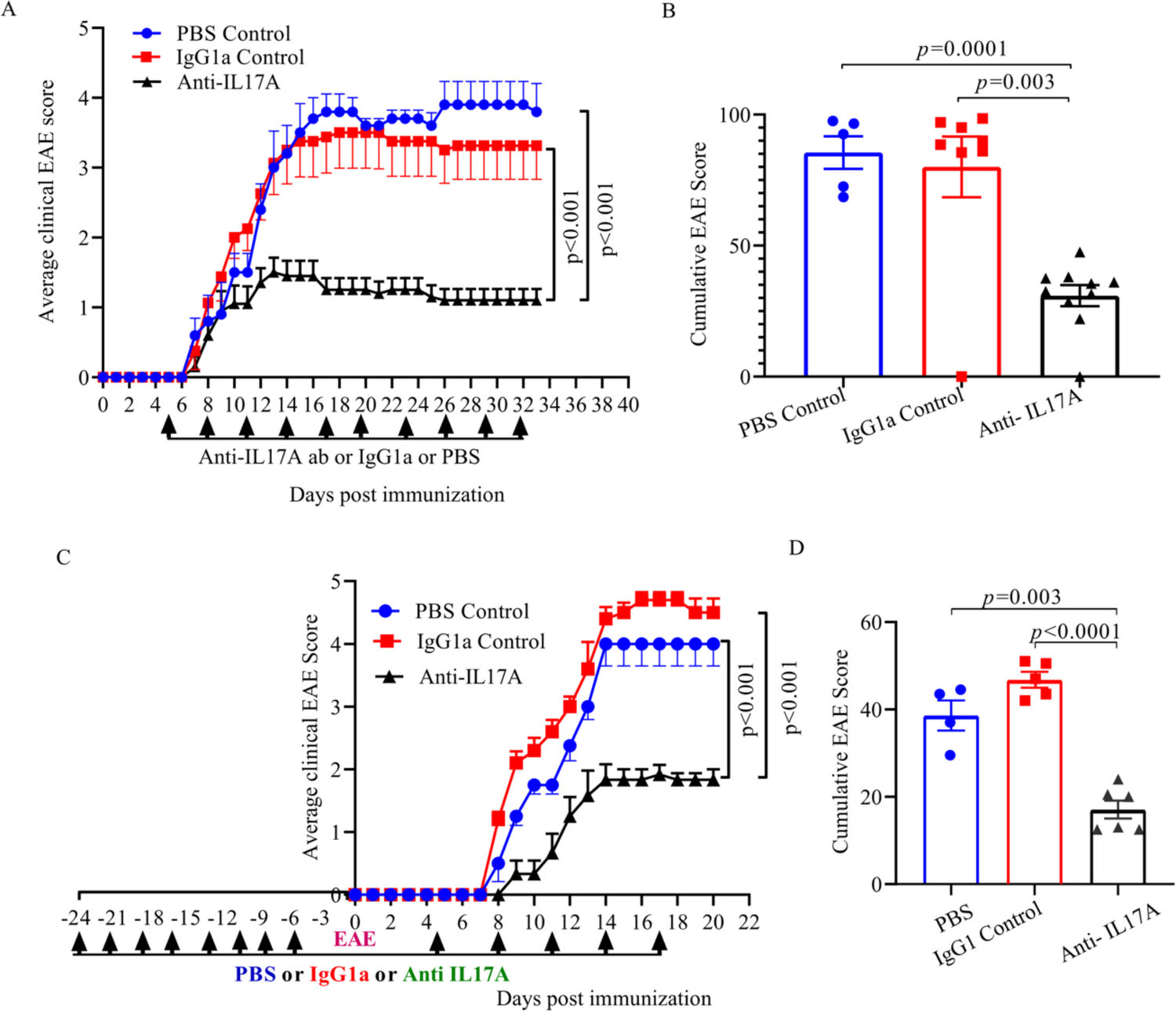
Anti-IL-17A treatment lowers the severity of MOG_35-55_ induced EAE in C57BL/6J mice. **(A-B)** 8-10-week-old B6 mice were immunized with MOG_35-55_/CFA. Five days after immunization, mice received anti-mouse IL-17A monoclonal antibody, isotype control mouse IgG1, or PBS intraperitoneally every three days until the end of the experiment (day 32). (A) Average clinical EAE scores over time. Treatment days are indicated by arrowheads. (B) Average cumulative EAE scores. (**C-D**) 8-10-week-old B6 mice were treated every three days with anti-IL-17A, IgG1, or PBS pre- and post-immunization with MOG_35-55_/CFA. (C) Average clinical EAE scores. Treatment days are indicated by arrowheads. (D) Average cumulative EAE scores. For plots in A, C, each bullet point represents the mean and bars represent the standard error of the mean from each experimental group. *p*-value was determined using two-way ANOVA following Tukey correction for EAE clinical scores (A, C) and unpaired t-test with Welch correction for cumulative EAE score (B, D). For graphs in B, D, each bar represents mean±SEM.

**Fig. S3:**
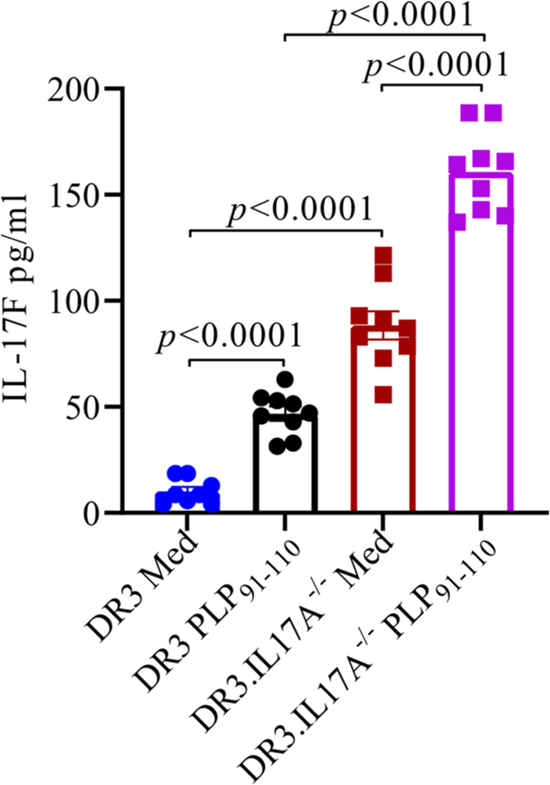
Levels of inflammatory cytokines IL-17F in DR3 and DR3.IL17A^-/-^ mice. 8-10-week-old mice were immunized with PLP_91-110_ /CFA to induce EAE. After 20 days, mice were euthanized, spleens were harvested, and splenic cells were cultured in the presence of PLP_91-110_ antigen for five days following stimulation with Phorbol myristate acetate (PMA) and ionomycin in the presence of Brefeldin A (BFA). The supernatant was collected and levels of IL17F were measured using ELISA. Cytokine values from DR3 mice were compared to those of DR3.IL17A^-/-^ mice in either media (Med) or after stimulation with PLP_91-110_. *p*-value was determined using unpaired t-test with Welch correction. Bars represent mean±S.E.M.

**Fig. S4:**
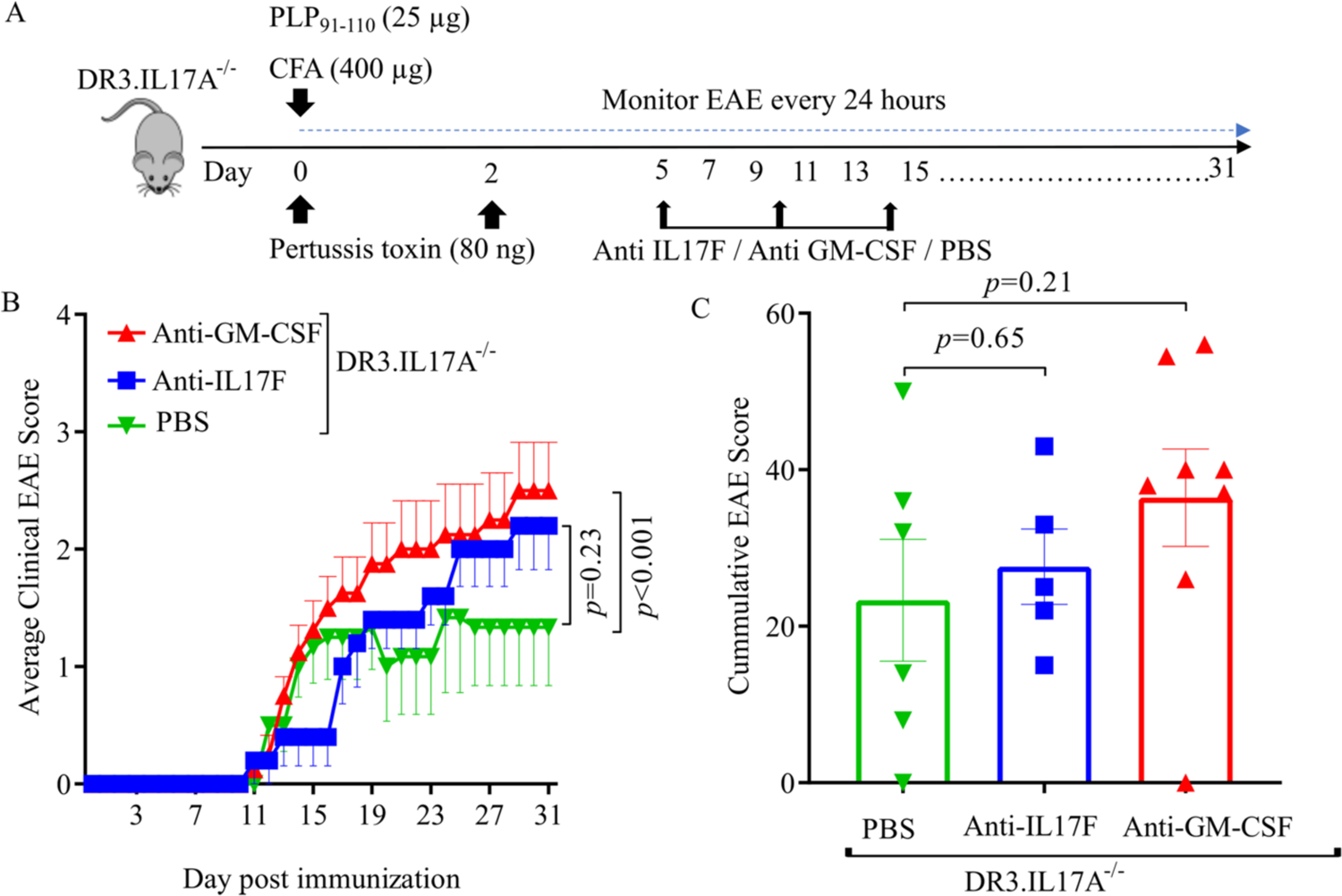
Treatment with anti-GM-CSF or anti-IL-17F monoclonal antibodies increases disease severity in DR3.IL17A^-/-^ mice. (**A**) Schematic diagram of the experimental workflow. EAE was induced in 8-10-week-old in DR3.IL17A^-/-^ transgenic mice. At days 5, 10, and 14 post-EAE induction, mice received intraperitoneal injections of either: 1) anti-GM-CSF monoclonal antibody, 2) anti-IL-17F monoclonal antibody, or 3) IgGa1 isotype control and clinical EAE scores were monitored for 31 days. (**B**) Average clinical EAE score of mice in each group over time. Bullet points represent the mean and bars represent the standard error of the mean of each experimental group from two independent experiments (n ≥3 mice per group). *p*-value was determined by a two-way ANOVA following Tukey correction for average clinical EAE score (B). (**C**) Average cumulative EAE scores. *p*-value was determined by unpaired t-test with Welch correction (C). For plots in B, C, each symbol represents the mean and bars represent the standard error of the mean from each experimental group.

**Fig. S5:**
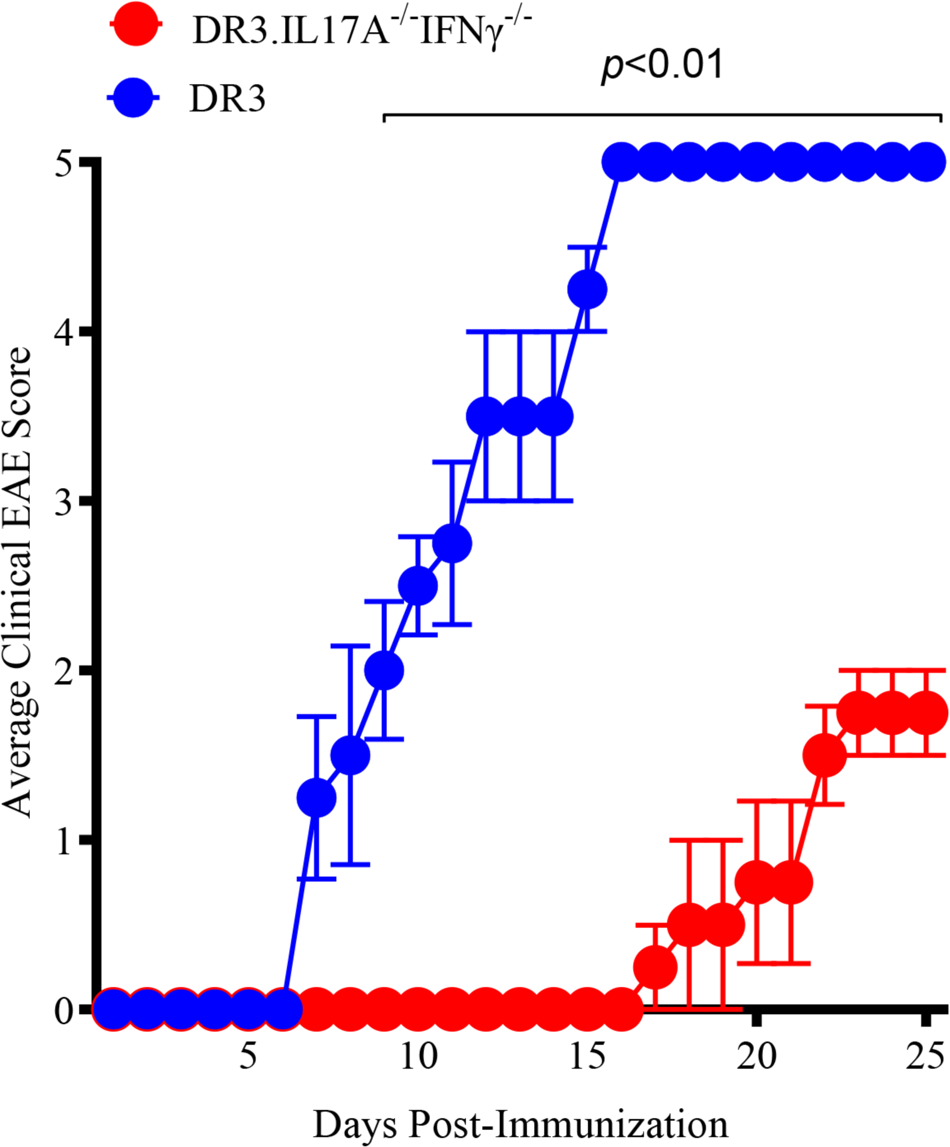
DR3.IL17A^-/-^ mice develops milder EAE in absence of IFNγ. EAE was induced in 8-10-week-old in DR3 and DR3.IL17A^-/-^IFNγ^-/-^ transgenic mice using PLP_91-110_, CFA and PTX. The disease was monitored and scored daily until day 25 of the experiment. Average clinical EAE score of mice in each group over time. Bullet points represent the mean and bars represent the standard error of the mean of each experimental group from two independent experiments (n =4 mice per group). *p*-value was determined by multiple unpaired t-tests with correction using two-stage step up method of Benjamini, Krieger and Yekutieli for average clinical EAE score.

**Fig. S6:**
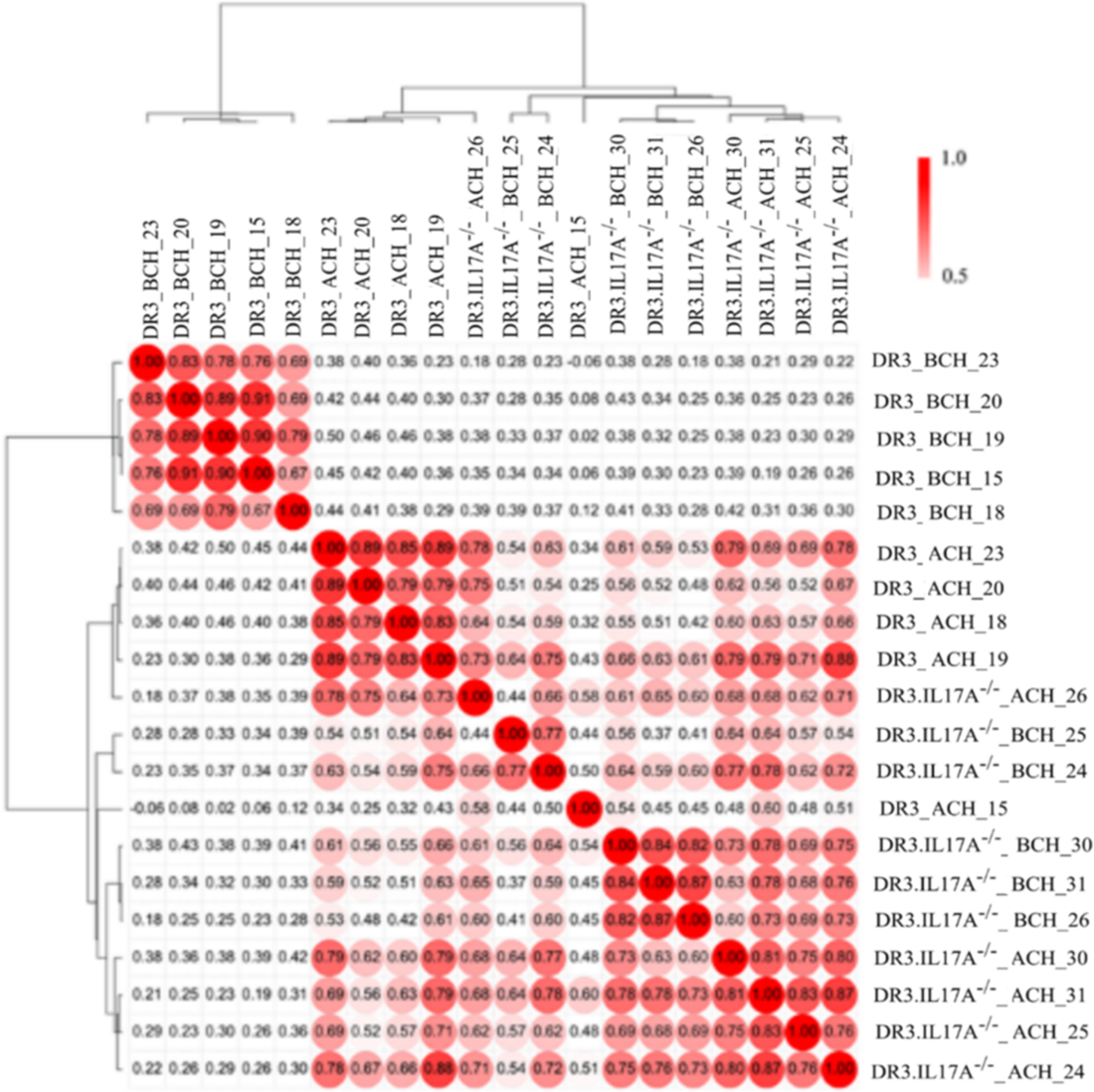
Similarity matrix of microbial compositional communities present in DR3 and DR3.IL17A^-/-^ mice before (BCH) and after (ACH) cohousing. Matrix was computed from the relative abundance matrix using 1-Spearman rank correlation metric using average linkage method and clustered hierarchically using Morpheus webserver (https://software.broadinstitute.org/morpheus).

**Fig. S7:**
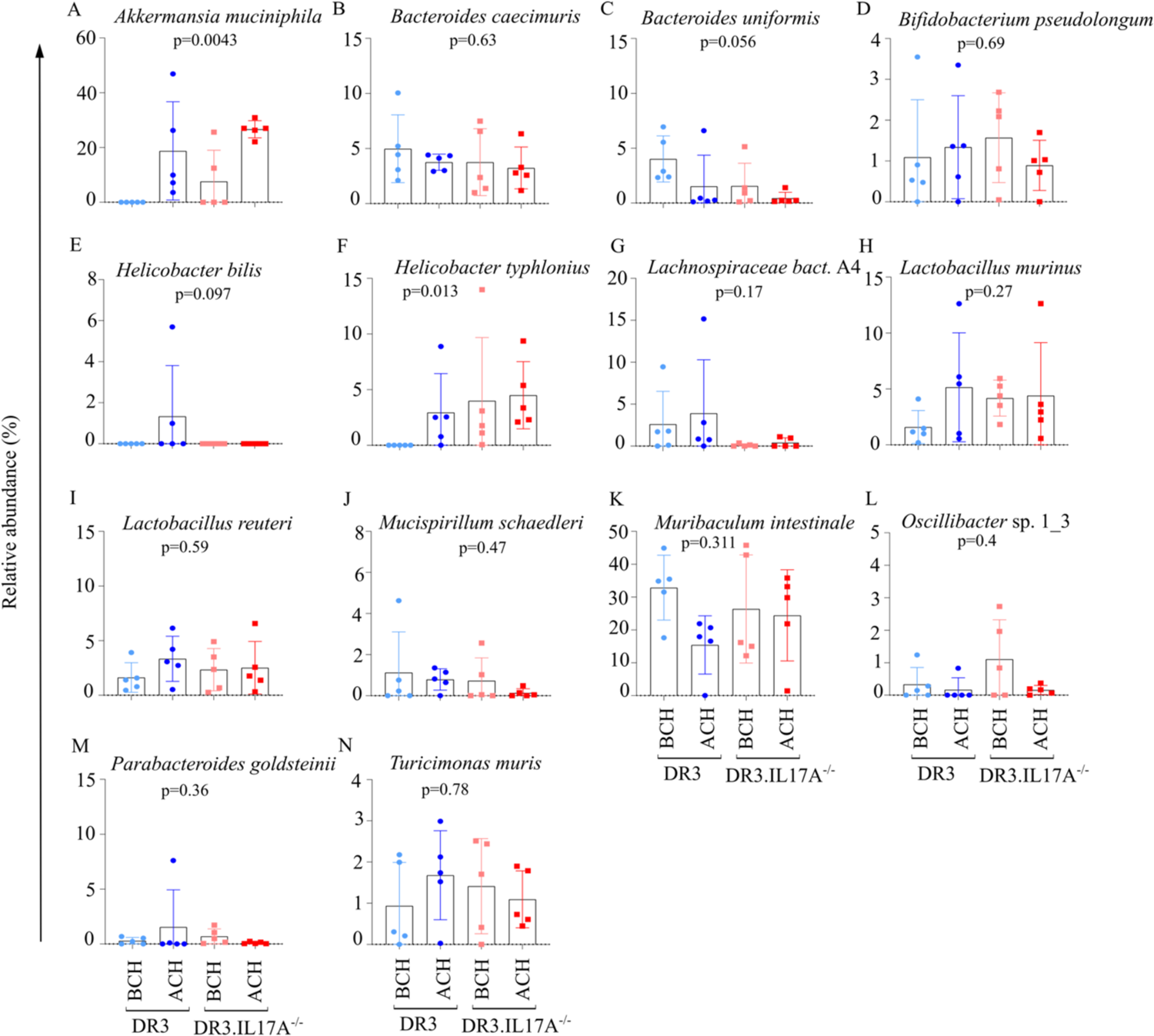
Composition of the gut bacteria in DR3 and DR3.IL17A^-/-^ transgenic mice before (BCH) and after (ACH) mice were cohoused. Each bar represents the average relative abundance (%) of the bacterium in each group. Kruskal Wallis test followed by Dunn-test was performed to identify the significance between the groups. ‘p-value’ from A-N represents the significance for the Kruskal Wallis test. Those bacteria with significant/non-significant p-value from Kruskal Wallis test but non-significantly different in at least one of the groups of interest (DR3-BCH to DR3.IL17A^-/-^-BCH, DR3-BCH to DR3-ACH and DR3.IL17A^-/-^-BCH to DR3.IL17A^-/-^-ACH) following Dunn test are shown.

**Fig. S8:**
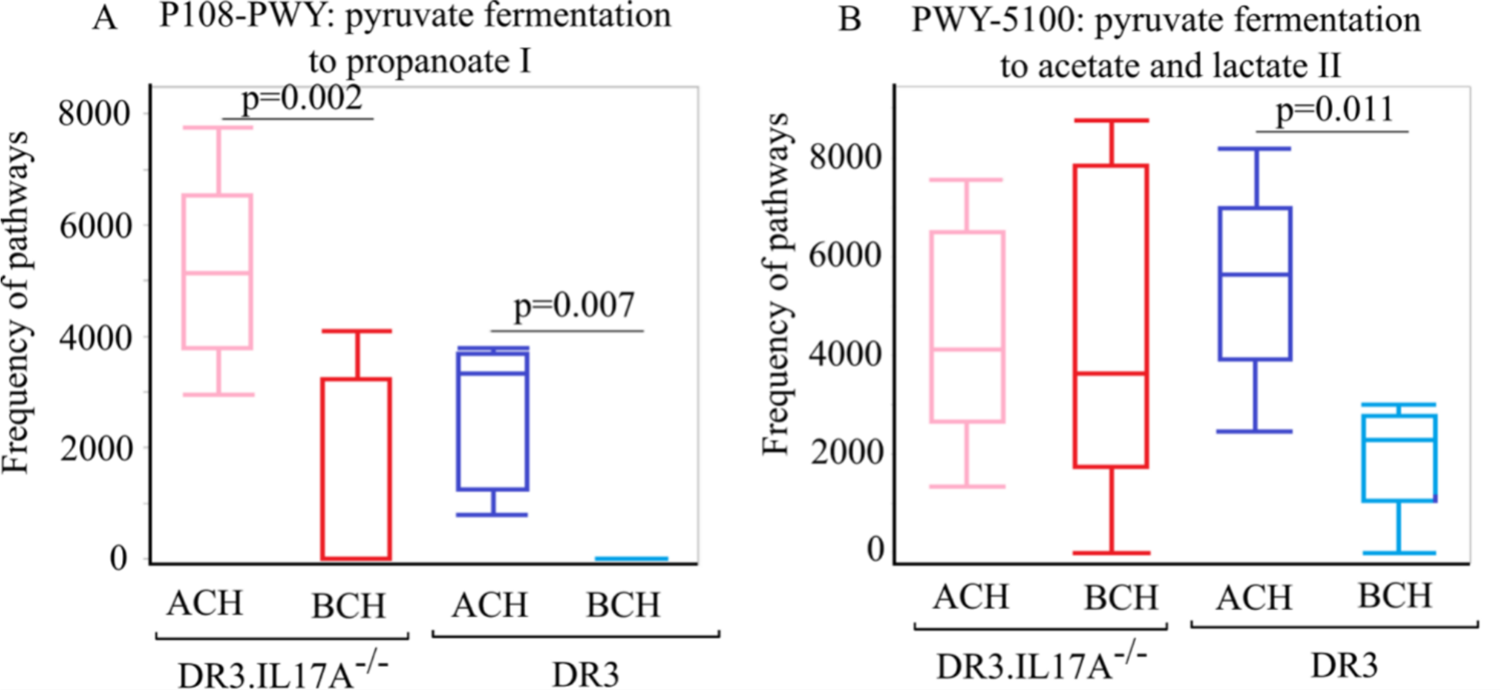
Abundance of short chain fatty acid pathways, as determined by shotgun metagenomics analysis in DR3.IL17A^-/-^ and DR3 transgenic mice before (BCH) and after (ACH) they were cohoused. Pairwise Wilcoxon test was performed to obtain the statistical significance BCH and ACH for each DR3 and DR3.IL17A^-/-^ transgenic mice.

**Fig. S9:**
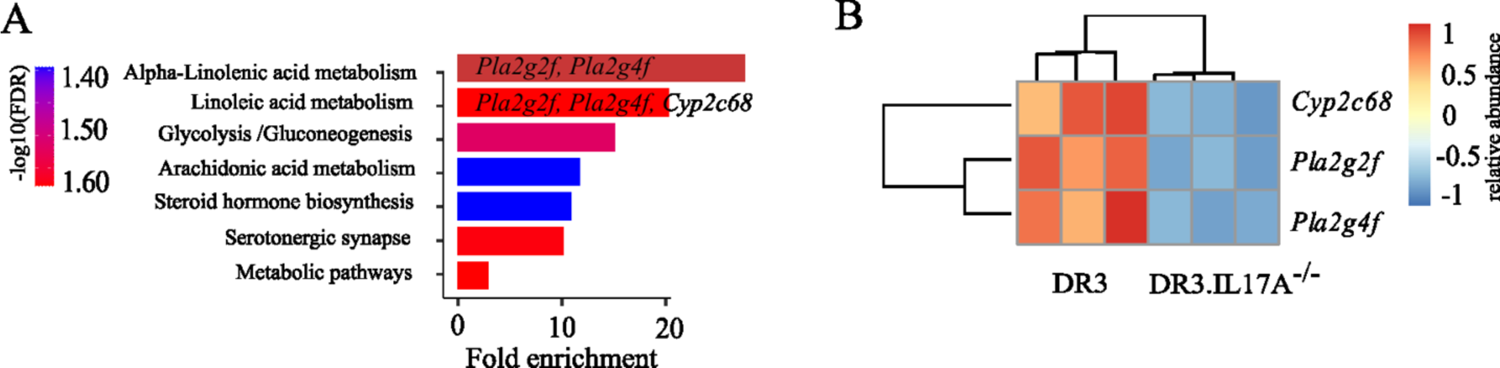
DR3 and DR3.IL17A^-/-^ transgenic mice exhibit distinct gene expression patterns in the colon. Colonic tissue was harvested from DR3 (n=3) or DR3.IL-17A^-/-^ transgenic mice (n=3) and subjected to RNA-seq analysis. A) Pathways enriched for 90 downregulated genes in KEGG database identified using ShinyGO webserver (37). B) Heatmap of enriched genes in alpha-linolenic and linoleic acid metabolism pathway. F) Expression of genes related to alpha-linolenic and linoleic acid metabolism pathway in DR3 or DR3.IL-17A^-/-^ transgenic mice.

---

**Table S1:**
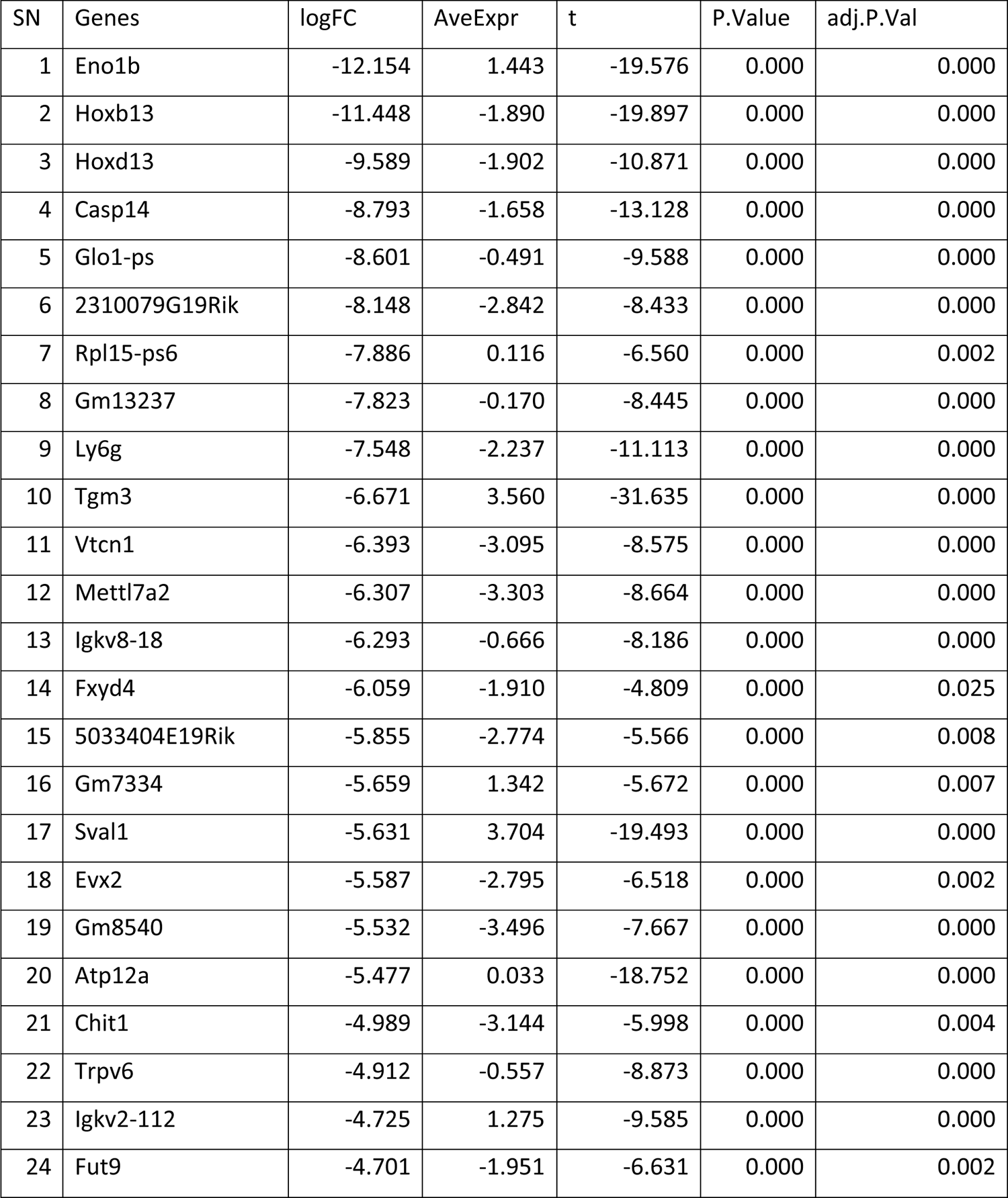
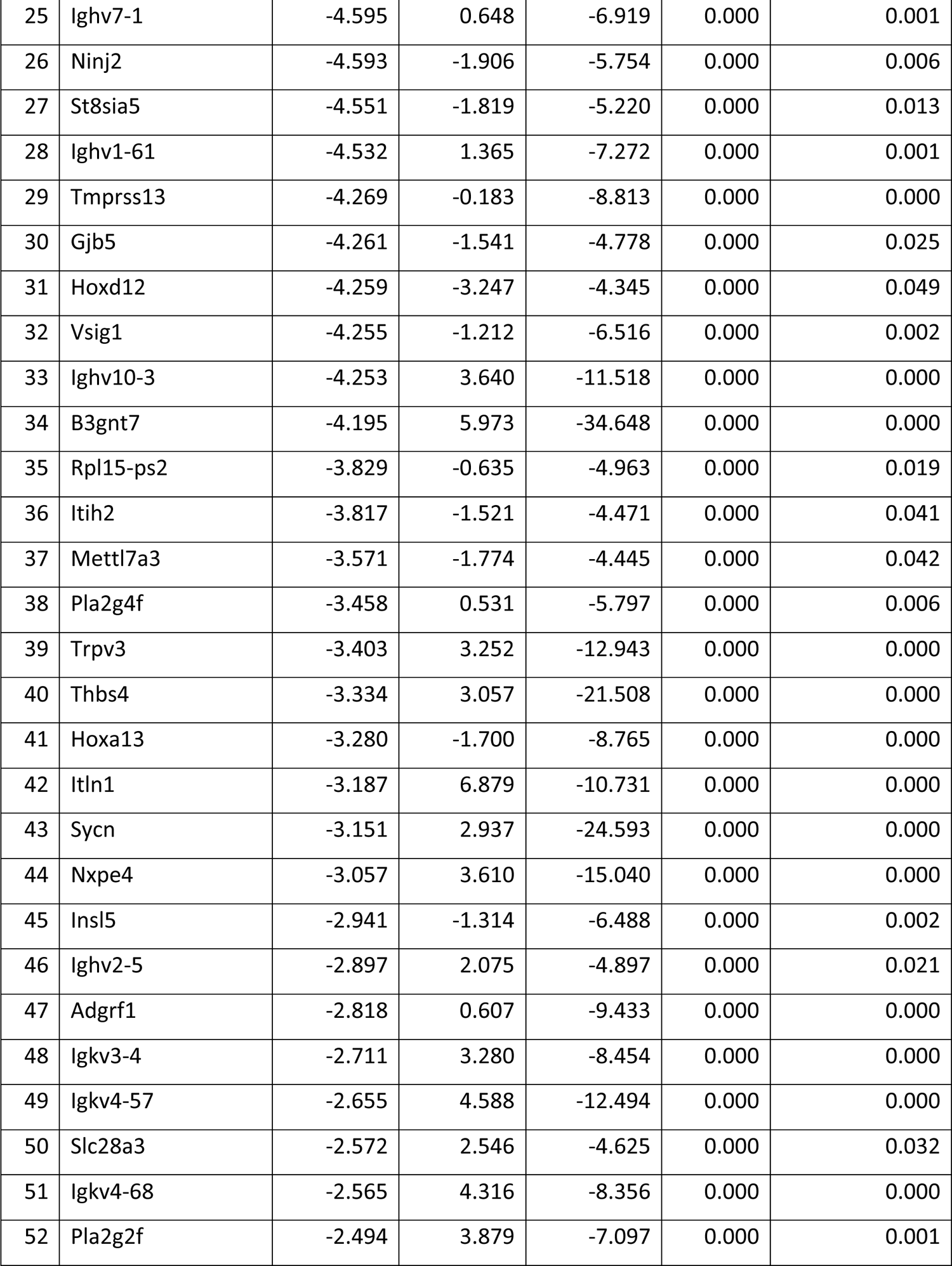
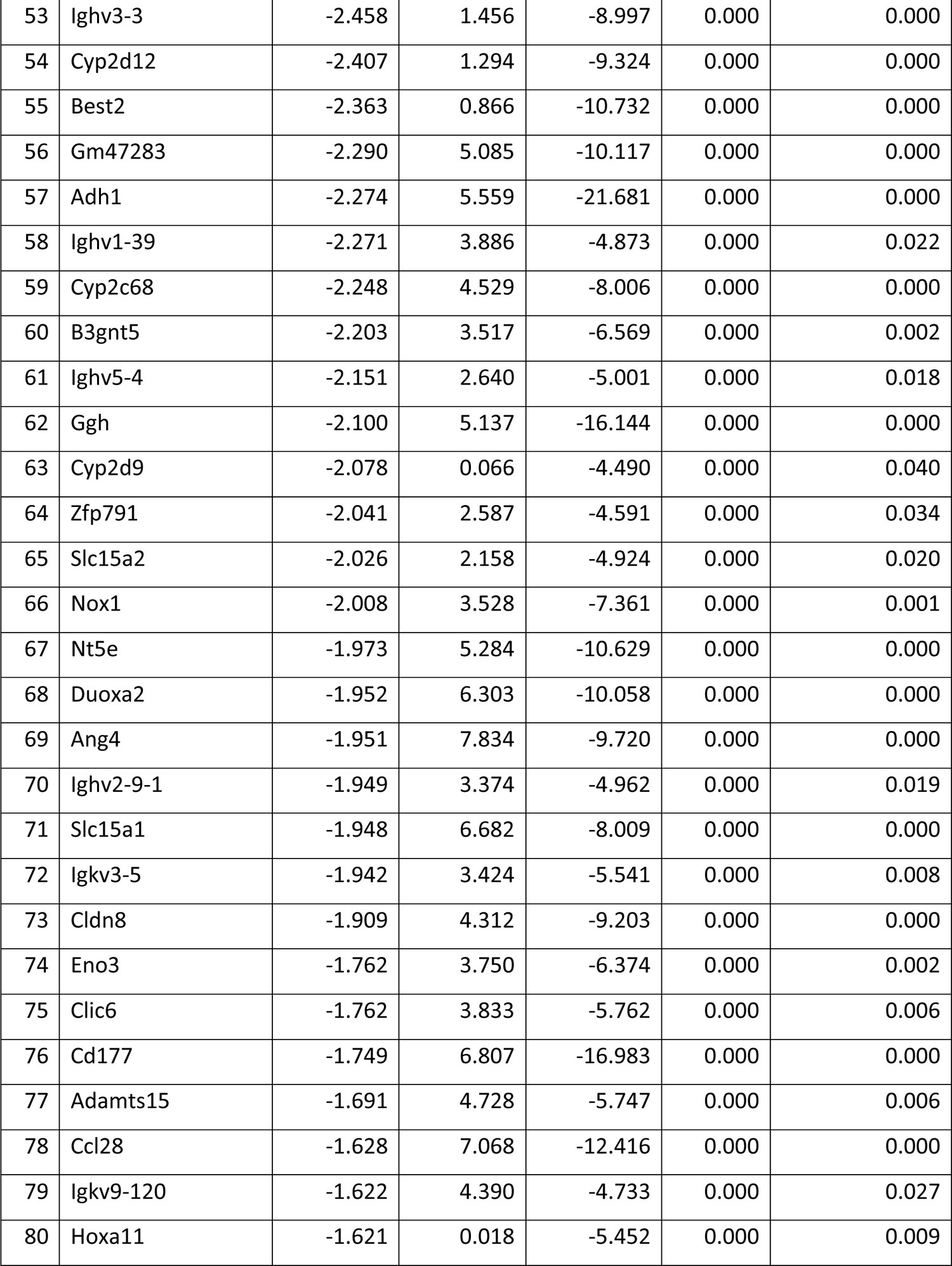
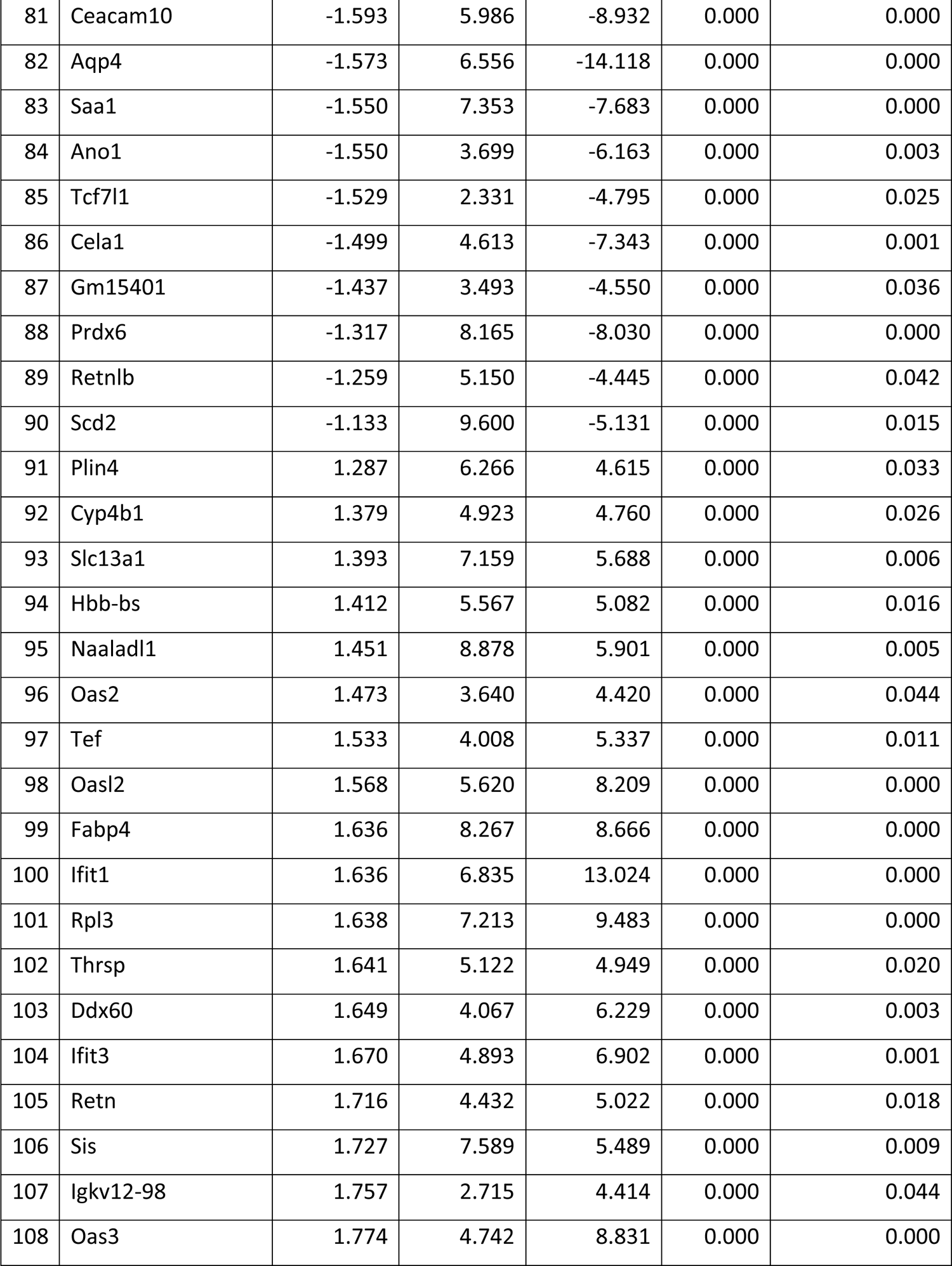
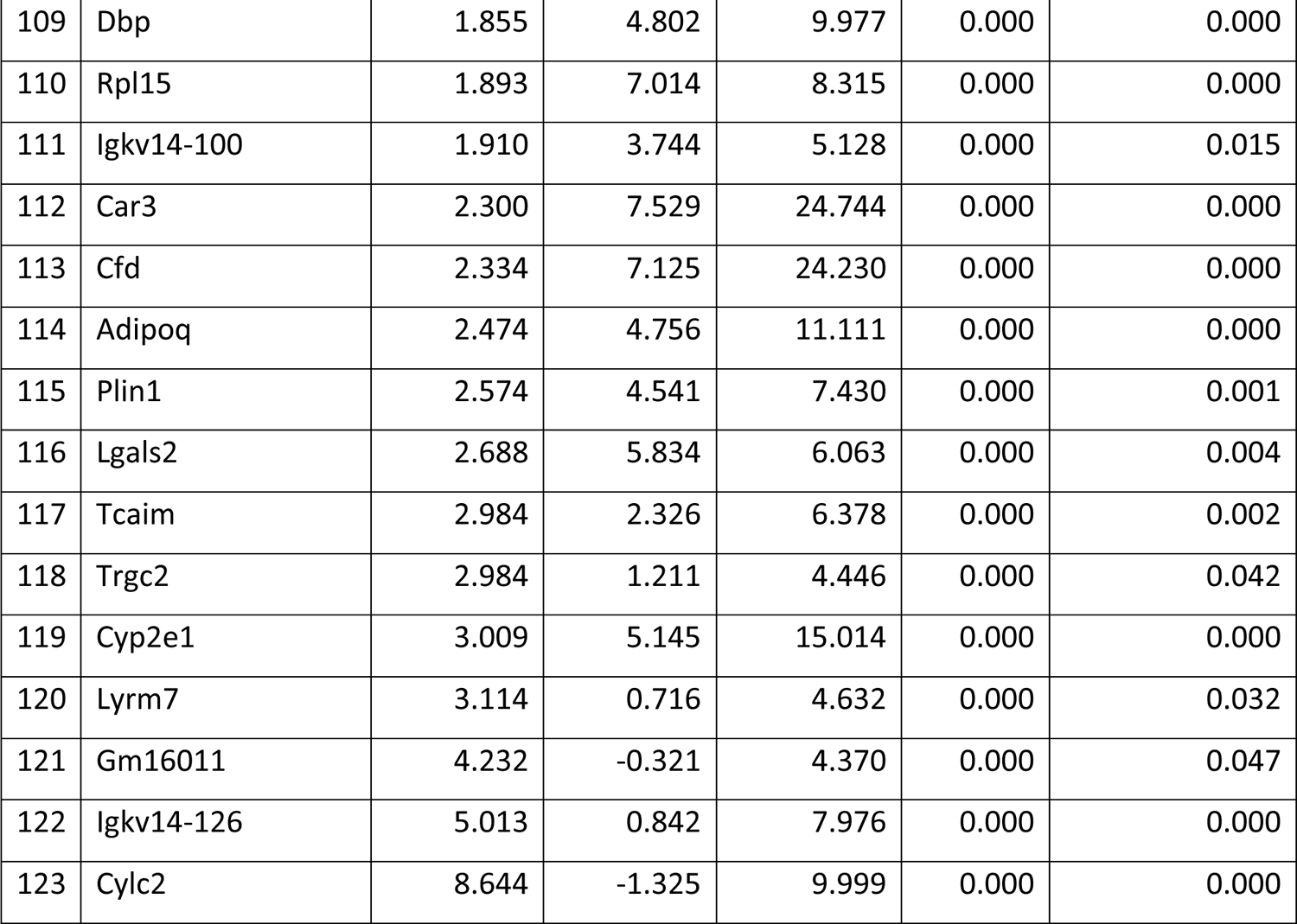
Differentially expressed gene list in the colon of DR3.IL17A^-/-^ compared to DR3 transgenic mice calculated using *limma* package within *edgeR* in R. All 123 differentially expressed genes with *LogFC*>1.0 and *adj.p.val*<0.05 are shown.

## References

1. Korn T, Bettelli E, Oukka M, Kuchroo VK. IL-17 and Th17 Cells. Annu Rev Immunol. 2009;27:485–517.

2. Milovanovic J, Arsenijevic A, Stojanovic B, Kanjevac T, Arsenijevic D, Radosavljevic G, et al. Interleukin-17 in Chronic Inflammatory Neurological Diseases. Front Immunol. 2020;11:947.

3. Curtis MM, Way SS. Interleukin-17 in host defence against bacterial, mycobacterial and fungal pathogens. Immunology. 2009;126(2):177–85.

4. Chong WP, Mattapallil MJ, Raychaudhuri K, Bing SJ, Wu S, Zhong Y, et al. The Cytokine IL-17A Limits Th17 Pathogenicity via a Negative Feedback Loop Driven by Autocrine Induction of IL-24. Immunity. 2020;53(2):384–97 e5.

5. Biswas PS. IL-17 in Renal Immunity and Autoimmunity. J Immunol. 2018;201(11):3153–9.

6. Iwakura Y, Nakae S, Saijo S, Ishigame H. The roles of IL-17A in inflammatory immune responses and host defense against pathogens. Immunol Rev. 2008;226:57–79.

7. McGeachy MJ, Cua DJ, Gaffen SL. The IL-17 Family of Cytokines in Health and Disease. Immunity. 2019;50(4):892–906.

8. Segal BM. Stage-specific immune dysregulation in multiple sclerosis. J Interferon Cytokine Res. 2014;34(8):633–40.

9. Weaver CT, Hatton RD. Interplay between the TH17 and TReg cell lineages: a (co-)evolutionary perspective. Nat Rev Immunol. 2009;9(12):883–9.

10. Yang Y, Torchinsky MB, Gobert M, Xiong H, Xu M, Linehan JL, et al. Focused specificity of intestinal TH17 cells towards commensal bacterial antigens. Nature. 2014;510(7503):152-6.

11. Atarashi K, Tanoue T, Oshima K, Suda W, Nagano Y, Nishikawa H, et al. Treg induction by a rationally selected mixture of Clostridia strains from the human microbiota. Nature. 2013;500(7461):232-6.

12. Alvarado-de la Barrera C, Zuniga-Ramos J, Ruiz-Morales JA, Estanol B, Granados J, Llorente L. HLA class II genotypes in Mexican Mestizos with familial and nonfamilial multiple sclerosis. Neurology. 2000;55(12):1897–900.

13. DeLuca GC, Ramagopalan SV, Herrera BM, Dyment DA, Lincoln MR, Montpetit A, et al. An extremes of outcome strategy provides evidence that multiple sclerosis severity is determined by alleles at the HLA-DRB1 locus. Proceedings of the National Academy of Sciences of the United States of America. 2007;104(52):20896–901.

14. Dyment DA, Herrera BM, Cader MZ, Willer CJ, Lincoln MR, Sadovnick AD, et al. Complex interactions among MHC haplotypes in multiple sclerosis: susceptibility and resistance. Hum Mol Genet. 2005;14(14):2019–26.

15. Olerup O, Hillert J. HLA class II-associated genetic susceptibility in multiple sclerosis: a critical evaluation. Tissue antigens. 1991;38(1):1–15.

16. Ramagopalan SV, Deluca GC, Morrison KM, Herrera BM, Dyment DA, Lincoln MR, et al. Analysis of 45 candidate genes for disease modifying activity in multiple sclerosis. Journal of neurology. 2008;255(8):1215–9.

17. Kaushansky N, Ben-Nun A. DQB1*06:02-Associated Pathogenic Anti-Myelin Autoimmunity in Multiple Sclerosis-Like Disease: Potential Function of DQB1*06:02 as a Disease-Predisposing Allele. Front Oncol. 2014;4:280.

18. Klehmet J, Shive C, Guardia-Wolff R, Petersen I, Spack EG, Boehm BO, et al. T cell epitope spreading to myelin oligodendrocyte glycoprotein in HLA-DR4 transgenic mice during experimental autoimmune encephalomyelitis. Clin Immunol. 2004;111(1):53–60.

19. Luckey D, Bastakoty D, Mangalam AK. Role of HLA class II genes in susceptibility and resistance to multiple sclerosis: studies using HLA transgenic mice. J Autoimmun. 2011;37(2):122–8.

20. Nalawade SA, Ji N, Raphael I, Pratt A, 3rd, Kraig E, Forsthuber TG. Aire is not essential for regulating neuroinflammatory disease in mice transgenic for human autoimmune-diseases associated MHC class II genes HLA-DR2b and HLA-DR4. Cell Immunol. 2018;331:38–48.

21. Quandt JA, Huh J, Baig M, Yao K, Ito N, Bryant M, et al. Myelin basic protein-specific TCR/HLA-DRB5*01:01 transgenic mice support the etiologic role of DRB5*01:01 in multiple sclerosis. J Immunol. 2012;189(6):2897–908.

22. Mangalam AK, Luo N, Luckey D, Papke L, Hubbard A, Wussow A, et al. Absence of IFN-gamma increases brain pathology in experimental autoimmune encephalomyelitis-susceptible DRB1*0301.DQ8 HLA transgenic mice through secretion of proinflammatory cytokine IL-17 and induction of pathogenic monocytes/microglia into the central nervous system. J Immunol. 2014;193(10):4859–70.

23. Mangalam A, Luckey D, Basal E, Behrens M, Rodriguez M, David C. HLA-DQ6 (DQB1*0601)-restricted T cells protect against experimental autoimmune encephalomyelitis in HLA-DR3.DQ6 double-transgenic mice by generating anti-inflammatory IFN-gamma. J Immunol. 2008; 180(11):7747-56.

24. Shahi SK, Freedman SN, Dahl RA, Karandikar NJ, Mangalam AK. Scoring disease in an animal model of multiple sclerosis using a novel infrared-based automated activity-monitoring system. Sci Rep. 2019;9(1):19194.

25. Tyler AF, Mendoza JP, Firan M, Karandikar NJ. CD8(+) T Cells Are Required For Glatiramer Acetate Therapy in Autoimmune Demyelinating Disease. PLoS One. 2013;8(6):e66772.

26. Mangalam A, Luckey D, Basal E, Jackson M, Smart M, Rodriguez M, et al. HLA-DQ8 (DQB1*0302)-restricted Th17 cells exacerbate experimental autoimmune encephalomyelitis in HLA-DR3-transgenic mice. J Immunol. 2009;182(8):5131–9.

27. Mangalam AK, Luckey D, Giri S, Smart M, Pease LR, Rodriguez M, et al. Two discreet subsets of CD8 T cells modulate PLP(91-110) induced experimental autoimmune encephalomyelitis in HLA-DR3 transgenic mice. J Autoimmun. 2012;38(4):344–53.

28. Goschl L, Preglej T, Hamminger P, Bonelli M, Andersen L, Boucheron N, et al. A T cell-specific deletion of HDAC1 protects against experimental autoimmune encephalomyelitis. J Autoimmun. 2018;86:51–61.

29. Thornton AM, Shevach EM. CD4+CD25+ immunoregulatory T cells suppress polyclonal T cell activation in vitro by inhibiting interleukin 2 production. J Exp Med. 1998;188(2):287–96.

30. Langmead B, Salzberg SL. Fast gapped-read alignment with Bowtie 2. Nat Methods. 2012;9(4):357–9.

31. Li D, Liu CM, Luo R, Sadakane K, Lam TW. MEGAHIT: an ultra-fast single-node solution for large and complex metagenomics assembly via succinct de Bruijn graph. Bioinformatics. 2015;31(10):1674–6.

32. Seemann T. Prokka: rapid prokaryotic genome annotation. Bioinformatics. 2014;30(14):2068–9.

33. Mangalam A, Shahi SK, Luckey D, Karau M, Marietta E, Luo N, et al. Human Gut-Derived Commensal Bacteria Suppress CNS Inflammatory and Demyelinating Disease. Cell Rep. 2017;20(6):1269–77.

34. Bray NL, Pimentel H, Melsted P, Pachter L. Near-optimal probabilistic RNA-seq quantification. Nat Biotechnol. 2016;34(5):525–7.

35. R Core Team. R: A language and environment for statistical computing R Foundation for Statistical Computing, Vienna, Austria. 2020.

36. Robinson MD, McCarthy DJ, Smyth GK. edgeR: a Bioconductor package for differential expression analysis of digital gene expression data. Bioinformatics. 2010;26(1):139–40.

37. Ge SX, Jung D, Yao R. ShinyGO: a graphical gene-set enrichment tool for animals and plants. Bioinformatics. 2020;36(8):2628–9.

38. Jensen SN, Cady NM, Shahi SK, Peterson SR, Gupta A, Gibson-Corley KN, et al. Isoflavone diet ameliorates experimental autoimmune encephalomyelitis through modulation of gut bacteria depleted in patients with multiple sclerosis. Sci Adv. 2021;7(28).

39. Komiyama Y, Nakae S, Matsuki T, Nambu A, Ishigame H, Kakuta S, et al. IL-17 plays an important role in the development of experimental autoimmune encephalomyelitis. J Immunol. 2006;177(1):566–73.

40. Duncker PC, Stoolman JS, Huber AK, Segal BM. GM-CSF Promotes Chronic Disability in Experimental Autoimmune Encephalomyelitis by Altering the Composition of Central Nervous System-Infiltrating Cells, but Is Dispensable for Disease Induction. J Immunol. 2018;200(3):966–73.

41. Li Z, Li D, Tsun A, Li B. FOXP3+ regulatory T cells and their functional regulation. Cell Mol Immunol. 2015;12(5):558–65.

42. Hori S, Nomura T, Sakaguchi S. Control of regulatory T cell development by the transcription factor Foxp3. Science. 2003;299(5609):1057-61.

43. Christensen AD, Skov S, Kvist PH, Haase C. Depletion of regulatory T cells in a hapten-induced inflammation model results in prolonged and increased inflammation driven by T cells. Clin Exp Immunol. 2015;179(3):485–99.

44. Montgomery TL, Kunstner A, Kennedy JJ, Fang Q, Asarian L, Culp-Hill R, et al. Interactions between host genetics and gut microbiota determine susceptibility to CNS autoimmunity. Proceedings of the National Academy of Sciences of the United States of America. 2020;117(44):27516–27.

45. Ridaura VK, Faith JJ, Rey FE, Cheng J, Duncan AE, Kau AL, et al. Gut microbiota from twins discordant for obesity modulate metabolism in mice. Science. 2013;341(6150):1241214.

46. Haghikia A, Jorg S, Duscha A, Berg J, Manzel A, Waschbisch A, et al. Dietary Fatty Acids Directly Impact Central Nervous System Autoimmunity via the Small Intestine. Immunity. 2015;43(4):817–29.

47. Mizuno M, Noto D, Kaga N, Chiba A, Miyake S. The dual role of short fatty acid chains in the pathogenesis of autoimmune disease models. PLoS One. 2017;12(2):e0173032.

48. Atarashi K, Tanoue T, Shima T, Imaoka A, Kuwahara T, Momose Y, et al. Induction of colonic regulatory T cells by indigenous Clostridium species. Science. 2011;331(6015):337–41.

49. Round JL, Mazmanian SK. Inducible Foxp3+ regulatory T-cell development by a commensal bacterium of the intestinal microbiota. Proceedings of the National Academy of Sciences of the United States of America. 2010;107(27):12204–9.

50. Miao Y, Zhang C, Yang L, Zeng X, Hu Y, Xue X, et al. The activation of PPARgamma enhances Treg responses through up-regulating CD36/CPT1-mediated fatty acid oxidation and subsequent N-glycan branching of TbetaRII/IL-2Ralpha. Cell Commun Signal. 2022;20(1):48.

51. Gao Z, Xu X, Li Y, Sun K, Yang M, Zhang Q, et al. Mechanistic Insight into PPARgamma and Tregs in Atherosclerotic Immune Inflammation. Front Pharmacol. 2021;12:750078.

52. Guo X, Dang W, Li N, Wang Y, Sun D, Nian H, et al. PPAR-alpha Agonist Fenofibrate Ameliorates Sjogren Syndrome-Like Dacryoadenitis by Modulating Th1/Th17 and Treg Cell Responses in NOD Mice. Invest Ophthalmol Vis Sci. 2022;63(6):12.

53. Wei X, Yang Z, Rey FE, Ridaura VK, Davidson NO, Gordon JI, et al. Fatty acid synthase modulates intestinal barrier function through palmitoylation of mucin 2. Cell Host Microbe. 2012;11(2):140–52.

54. Shahi SK, Freedman SN, Murra AC, Zarei K, Sompallae R, Gibson-Corley KN, et al. Prevotella histicola, A Human Gut Commensal, Is as Potent as COPAXONE(R) in an Animal Model of Multiple Sclerosis. Front Immunol. 2019;10:462.

55. Regen T, Isaac S, Amorim A, Nunez NG, Hauptmann J, Shanmugavadivu A, et al. IL-17 controls central nervous system autoimmunity through the intestinal microbiome. Sci Immunol. 2021;6(56).

56. McGinley AM, Sutton CE, Edwards SC, Leane CM, DeCourcey J, Teijeiro A, et al. Interleukin-17A Serves a Priming Role in Autoimmunity by Recruiting IL-1beta-Producing Myeloid Cells that Promote Pathogenic T Cells. Immunity. 2020;52(2):342–56 e6.

57. Ghosh D, Curtis AD, 2nd, Wilkinson DS, Mannie MD. Depletion of CD4+ CD25+ regulatory T cells confers susceptibility to experimental autoimmune encephalomyelitis (EAE) in GM-CSF-deficient Csf2-/- mice. J Leukoc Biol. 2016;100(4):747–60.

58. Haak S, Croxford AL, Kreymborg K, Heppner FL, Pouly S, Becher B, et al. IL-17A and IL-17F do not contribute vitally to autoimmune neuro-inflammation in mice. J Clin Invest. 2009;119(1):61–9.

59. Wanke F, Tang Y, Gronke K, Klebow S, Moos S, Hauptmann J, et al. Expression of IL-17F is associated with non-pathogenic Th17 cells. J Mol Med (Berl). 2018;96(8):819–29.

60. McQualter JL, Darwiche R, Ewing C, Onuki M, Kay TW, Hamilton JA, et al. Granulocyte macrophage colony-stimulating factor: a new putative therapeutic target in multiple sclerosis. J Exp Med. 2001;194(7):873–82.

61. Pierson ER, Goverman JM. GM-CSF is not essential for experimental autoimmune encephalomyelitis but promotes brain-targeted disease. JCI Insight. 2017;2(7):e92362.

62. Cekanaviciute E, Yoo BB, Runia TF, Debelius JW, Singh S, Nelson CA, et al. Gut bacteria from multiple sclerosis patients modulate human T cells and exacerbate symptoms in mouse models. Proc Natl Acad Sci U S A. 2017;114(40):10713–8.

63. Nutsch KM, Hsieh CS. T cell tolerance and immunity to commensal bacteria. Curr Opin Immunol. 2012;24(4):385–91.

64. Lozupone CA, Stombaugh JI, Gordon JI, Jansson JK, Knight R. Diversity, stability and resilience of the human gut microbiota. Nature. 2012;489(7415):220-30.

65. Sun M, Wu W, Chen L, Yang W, Huang X, Ma C, et al. Microbiota-derived short-chain fatty acids promote Th1 cell IL-10 production to maintain intestinal homeostasis. Nat Commun. 2018;9(1):3555.

66. Kashi VP, Ortega SB, Karandikar NJ. Neuroantigen-specific autoregulatory CD8+ T cells inhibit autoimmune demyelination through modulation of dendritic cell function. PLoS One. 2014;9(8):e105763.

67. Kaisar MMM, Pelgrom LR, van der Ham AJ, Yazdanbakhsh M, Everts B. Butyrate Conditions Human Dendritic Cells to Prime Type 1 Regulatory T Cells via both Histone Deacetylase Inhibition and G Protein-Coupled Receptor 109A Signaling. Front Immunol. 2017;8:1429.

68. Sohn JH, Lee YK, Han JS, Jeon YG, Kim JI, Choe SS, et al. Perilipin 1 (Plin1) deficiency promotes inflammatory responses in lean adipose tissue through lipid dysregulation. J Biol Chem. 2018;293(36):13974–88.

69. Chwojnicki K, Iwaszkiewicz-Grzes D, Jankowska A, Zielinski M, Lowiec P, Gliwinski M, et al. Administration of CD4(+)CD25(high)CD127(-)FoxP3(+) Regulatory T Cells for Relapsing-Remitting Multiple Sclerosis: A Phase 1 Study. BioDrugs. 2021;35(1):47–60.

